# The evolution of sex-biased gene expression in the *Drosophila* brain

**DOI:** 10.1101/2020.04.14.041624

**Authors:** Samuel Khodursky, Nicolas Svetec, Sylvia Durkin, Li Zhao

## Abstract

Genes with sex-biased expression in *Drosophila* are thought to underlie sexually dimorphic phenotypes and have been shown to possess important evolutionary properties. However, the forces and constraints governing the evolution of sex-biased genes in the somatic tissues of *Drosophila* are largely unknown. Using population-scale RNA sequencing data we show that sex-biased genes in the *Drosophila* brain are highly enriched on the X Chromosome and that most are biased in a species-specific manner. We show that X-linked male-biased genes, and to a lesser extent female-biased genes, are enriched for signatures of directional selection at the gene expression level. By examining the evolutionary properties of gene flanking regions on the X Chromosome, we find evidence that adaptive cis-regulatory changes are more likely to drive the expression evolution of X-linked male-biased genes than other X-linked genes. Finally, we examine whether constraint due to broad expression across multiple tissues and genetic constraint due to the largely shared male and female genomes could be responsible for the observed patterns of gene expression evolution. We find that expression breadth does not constrain the directional evolution of gene expression in the brain. Additionally, we find that the shared genome between males and females imposes a substantial constraint on the expression evolution of sex-biased genes. Overall, these results significantly advance our understanding of the patterns and forces shaping the evolution of sexual dimorphism in the *Drosophila* brain.

## Introduction

Even though males and females share nearly identical genomes — with the exception of the Y Chromosome — they exhibit clear phenotypic differences at both the physiological, morphological, and behavioral levels (Darwin 1871; Arnqvist and Rowe 2005). Thus, one of the most exciting challenges in evolutionary biology is understanding how males and females use shared genomes to produce sexually dimorphic phenotypes. A popular and tractable approach to understanding sexual dimorphism is to examine differences between the sexes at the transcriptome level (Jin et al. 2001; Ellegren and Parsch 2007; Grath and Parsch 2016). Historically, most work in *Drosophila* has focused on genes with sex-biased expression in either gonad or whole body samples, where the differential expression signal is dominated by gonad expression (Ranz et al. 2003; Parisi et al. 2003; Whittle and Extavour 2019). Studies examining the forces influencing the evolution of these genes found that sex-biased genes — and male-biased genes in particular — are highly enriched for genes exhibiting signatures of positive selection at the protein level, suggesting that the evolution of sex-biased genes is central to adaptive evolution in *Drosophila* (Pröschel et al. 2006; Grath and Parsch 2016). Relative to female-biased and unbiased genes, the expression levels of male-biased genes tend to diverge rapidly between species (Zhang et al. 2007; Ranz et al. 2003). However, genes frequently gain and lose sex-bias over evolutionary time (Zhang et al. 2007; Ranz et al. 2003). It has also been shown that sex-biased genes within gonads and whole body samples are distributed among chromosomes in a highly non-random manner: male-biased genes are significantly depleted on the X Chromosome while female-biased genes are significantly enriched on the X Chromosome, which may be attributable to the lack of X Chromosome dosage compensation in the male germline (Meiklejohn et al. 2011; Meiklejohn and Presgraves 2012; Parisi et al. 2003). Finally, several studies have examined sex-biased gene expression in the *D. melanogaster* brain (Catalán et al. 2012; Huylmans and Parsch 2015; Pacifico et al. 2018). In contrast to findings in the gonads, male- and female-biased genes were both enriched on the X Chromosome (Catalán et al. 2012; Huylmans and Parsch 2015), suggesting that the chromosomal distributions of genes with sex-biased expression are highly tissue dependent. Despite these advances, the evolutionary properties of genes with sex-biased expression in sexually dimorphic somatic tissues such as the *Drosophila* brain remain largely unknown (Catalán et al. 2012; Chang et al. 2011).

Some possible forces and constraints governing expression evolution include the faster-X effect, expression breadth across tissues, and genetic constraint. The faster-X effect is a phenomenon where genes and regulatory regions tend to evolve more rapidly on the X Chromosome than on the autosomes (Charlesworth et al. 1987; Meisel et al. 2012; Meisel and Connallon 2013). This can be due to the relatively rapid fixation of beneficial recessive mutations on the X Chromosome, and this effect has been observed in both sex-biased and unbiased genes but is usually most prominent in male-biased genes (Charlesworth et al. 1987; Begun et al. 2007; Baines et al. 2008; Meisel et al. 2012; Meisel and Connallon 2013; Orr and Betancourt 2001; Vicoso and Charlesworth 2006). Broad expression across multiple tissues may also act to constrain the expression evolution of genes in the brain. For instance, genes which are expressed in many tissues are likely to have an important role in at least several tissues and hence their expression levels may be subjected to greater selective constraint (Orr 2000; Chen and Dokholyan 2006; Meisel et al. 2012; Papakostas et al. 2014). Additionally, the genetic constraints imposed by the largely shared male and female genomes have been shown to constrain the evolution of sexually dimorphic phenotypes in various species including *Drosophila* (Poissant et al. 2010; Griffin et al. 2013). By understanding the forces and constraints governing the evolution of sex-biased gene expression in the *Drosophila* brain, we may begin to uncover the molecular rules underpinning the evolution of sexually dimorphic behaviors.

In order to understand the evolutionary properties of genes exhibiting sex-biased expression in the brain, we generated and analyzed gene expression data from female and male brains in multiple strains of *D. melanogaster, D. simulans*, and *D. yakuba*. To determine the types of selection affecting expression levels, we examined interspecific and intraspecific variability in gene expression using an analysis of variance (ANOVA) framework. Genes with a large ratio of interspecific to intraspecific variability are likely evolving under directional selection. Genes with low interspecific and intraspecific variabilities are likely evolving under stabilizing selection. Low interspecific and high intraspecific variability is indicative of possible balancing selection or adaptive variation (Nuzhdin et al. 2004). We found that X-linked sex-biased genes are enriched for signatures of directional selection at the expression level, while autosomal female- and male-biased genes are enriched for signatures of balancing and stabilizing selection, respectively. Additionally, we discovered that broad expression across tissues does not constrain the directional evolution of gene expression in the *Drosophila* brain but does constrain expression variability. Finally, we showed that the shared genome between males and females imposes a significant constraint on the expression evolution of sex-biased genes in the *Drosophila* brain.

## Results

### Large enrichment of sex-biased genes on the X Chromosome

Using strand-specific poly-A+ RNA-seq data collected from the brains of *D. melanogaster, D. simulans*, and *D. yakuba* (**Figure 1A**), we analyzed the expression patterns of each species, and identified 1246 female- and 989 male-biased genes in *D. melanogaster*, 767 female- and 725 male-biased genes in *D. simulans*, and 417 female- and 551 male-biased genes in *D. yakuba* at a false-discovery rate (FDR) of 0.05 (**Figure 1B, Supplementary Table S1**). By looking at the chromosomal distribution of female- and male-biased genes, we found that both sets of genes were highly enriched on the X Chromosome in all three species (p<1×10^−5^, by a factor of about 2-3 fold in all species) and depleted on the autosomes, suggesting that the enrichment of X-linked genes with sex-biased expression in the brain is a conserved phenomenon (**Figure 2**).

**Figure 1.**
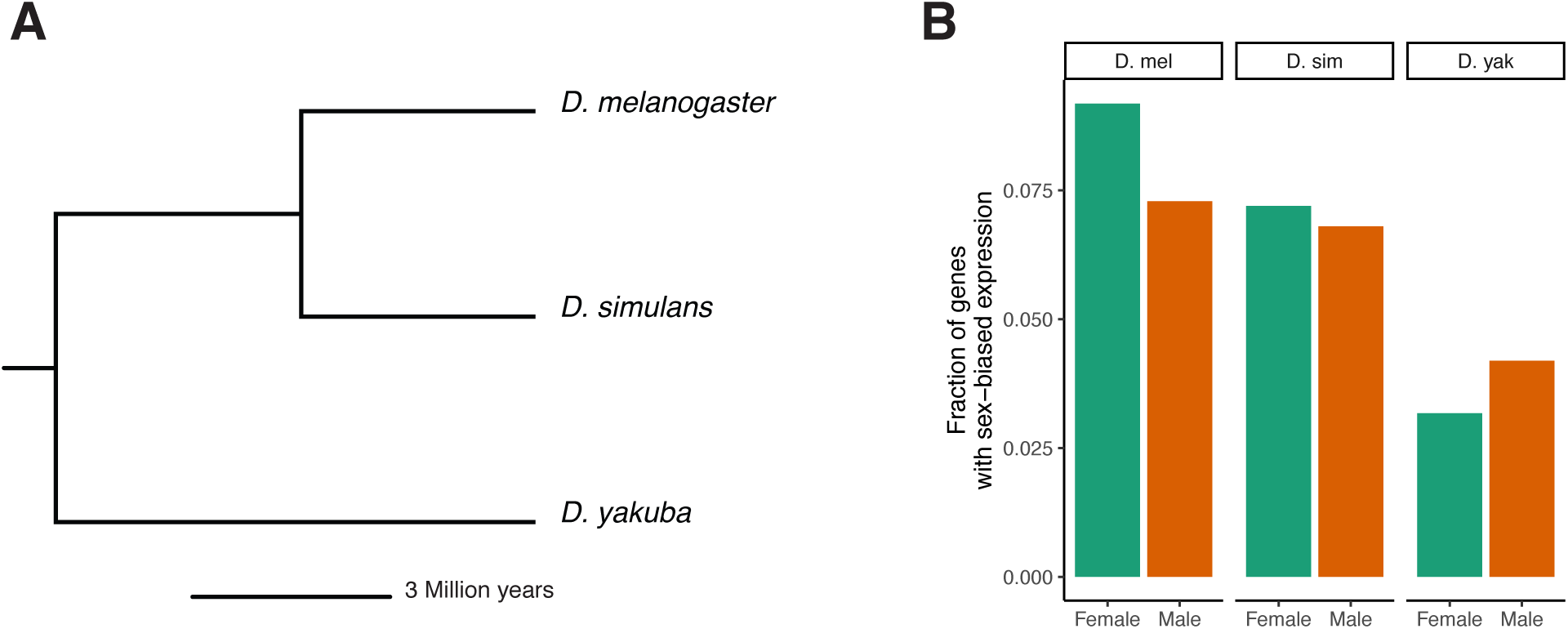
Species tree and number of sex-biased genes. A) A phylogenetic tree with the three species studied. B) The fraction of the expressing genome showing sex-biased expression in each of the three species at a FDR of 0.05. The expressing genome was defined to be the set of all genes with finite p-values (from DESeq2) for a given species.

**Figure 2.**
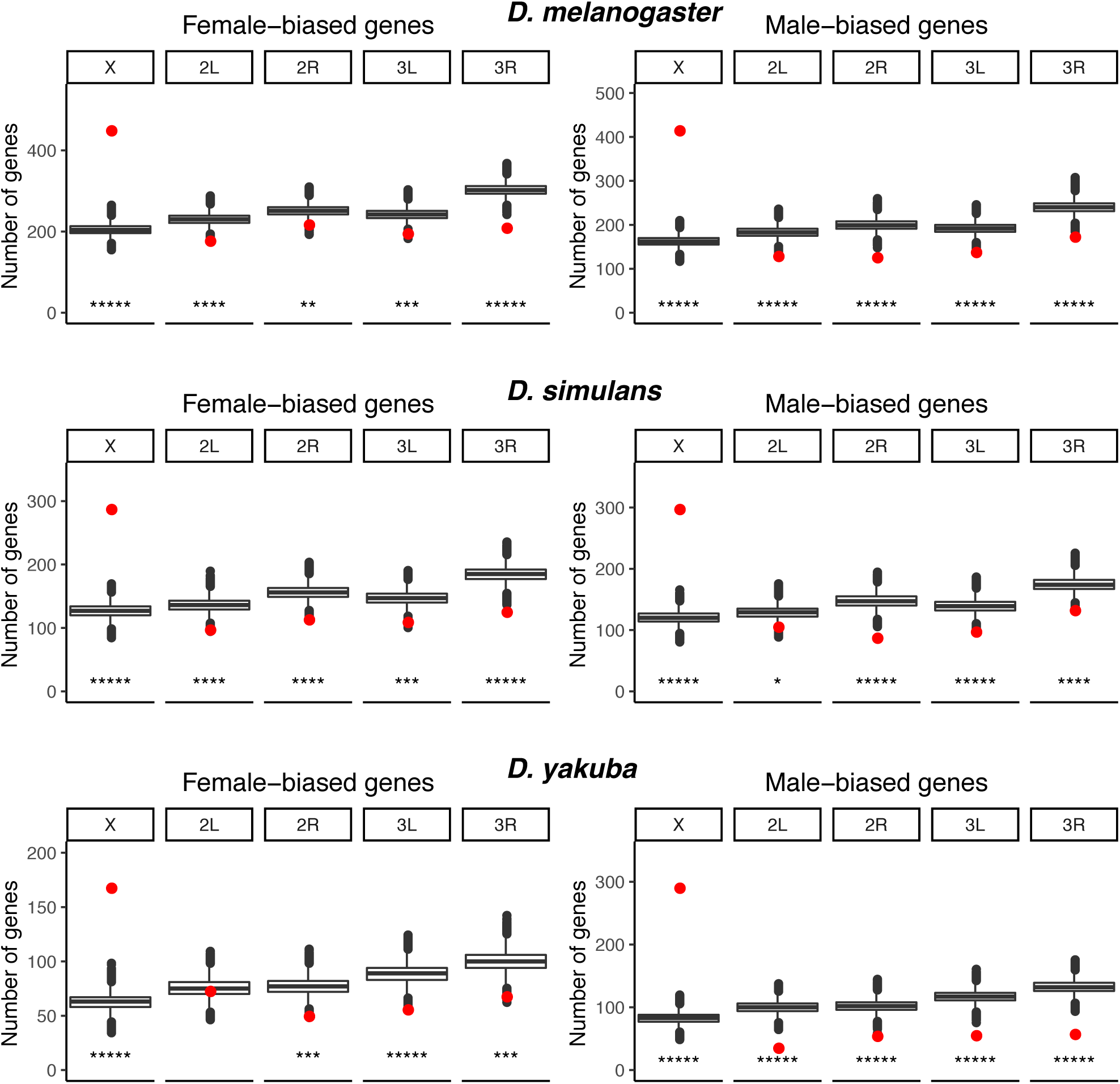
Chromosomal distributions of genes with sex-biased expression in the *Drosophila* brain. Sex-biased genes are significantly enriched on the X Chromosome and depleted on the autosomes in the three *Drosophila* species. The red points indicate the observed number of sex-biased genes on a given chromosome arm (at a FDR of 0.05) and the box plots indicate the simulated numbers of sex-biased genes. *****p-value < 1×10^−5^, ****p-value < 1×10^−4^, ***p-value < 1×10^−3^, **p-value < 1×10^−2^, *p-value < 5×10^−2^. The p-values shown are 2 times the fraction of simulations which resulted in a more extreme number of sex-biased genes on a given chromosome arm.

To determine if the significant enrichment of sex-biased genes on the X Chromosome was unique to the brain, we analyzed previously published expression data from *D. melanogaster* and *D. yakuba* in four other tissues: gonads, thorax, abdomen, and viscera (Yang et al. 2018). We found that the large enrichment of sex-biased genes on the X Chromosome was unusually large in the brain in both species (**Supplementary Figure S1**).

To determine if certain biological processes were enriched in the sets of sex-biased genes, we used PANTHER gene ontology (Carbon et al. 2019; Ashburner et al. 2000) on lists of X chromosomal and autosomal male- and female-biased genes in all three species at an FDR of 0.05. In all three species, male-biased autosomal genes were enriched for gene-ontology terms relating to metabolic processes (**Supplementary Table S2**). Other classes of genes were not consistently enriched for similar terms across the three species. In fact, with the exception of male-biased genes in *D. yakuba*, X-linked sex-biased genes were not enriched for any terms in any of the species.

We compared the expression levels of sex-biased and unbiased genes. The expression patterns were largely consistent between the three species, and male-biased autosomal genes had relatively high expression (**Supplementary Figure S2**). We then examined the expression fold-changes and significance-levels of sex-biased genes and found that in all three species X Chromosome male-biased genes had a significantly higher fold-change than autosomal male-biased genes (**Supplementary Figure S3**). Moreover, in all three species and in both sexes, sex-biased genes on the X Chromosome were more significantly sex-biased (smaller in q-value) than genes on the autosomes (**Supplementary Figure S3**). Given that female-biased X Chromosome genes did not show higher fold-changes than autosomal female-biased genes (with the exception of *D. simulans*), but were more significantly sex-biased, our results suggest that female-biased autosomal genes may exhibit a high degree of expression variability.

### Rates of sex-bias turnover are high and chromosome-specific

These observations prompted us to look at the evolution of sex-biased genes at the expression level. First, we looked at the turnover of sex-biased genes between the three *Drosophila* species. By only considering annotated 1:1:1 orthologs (from FlyBase (Thurmond et al. 2019)) we found that 57% of male-biased genes and 65% of female-biased genes within the three species are sex-biased in a species-specific manner (**Figure 3**), supporting that there is rapid sex-biased gene expression turnover in the brain. This result is consistent with findings in *Heliconius* butterflies (Catalán et al. 2018), suggesting that fast evolution and turnover of sex-biased genes in the brain may be a pervasive and conserved phenomenon in animals. In addition, we found the ratio of multi-species to single-species sex-biased genes to be substantially higher on the X Chromosome compared to the autosomes in both sexes (male-biased: 1.42 vs 0.46, female-biased: 1.01 vs 0.34, *p* < 2.2×10^−16^ for both, Fisher’s exact test), suggesting that chromosome-dependent patterns of selection at the level of gene expression may be responsible for the accumulation of sex-biased genes on the X Chromosome. We also observed that a large majority (69 out of 97, **Supplementary Table S3**) of sex-biased genes which have ‘switched’ sex-bias at some point in their recent evolutionary history (i.e. they are male-biased in at least one species and female-biased in at least one other *Drosophila* species) were located on the X Chromosome.

**Figure 3.**
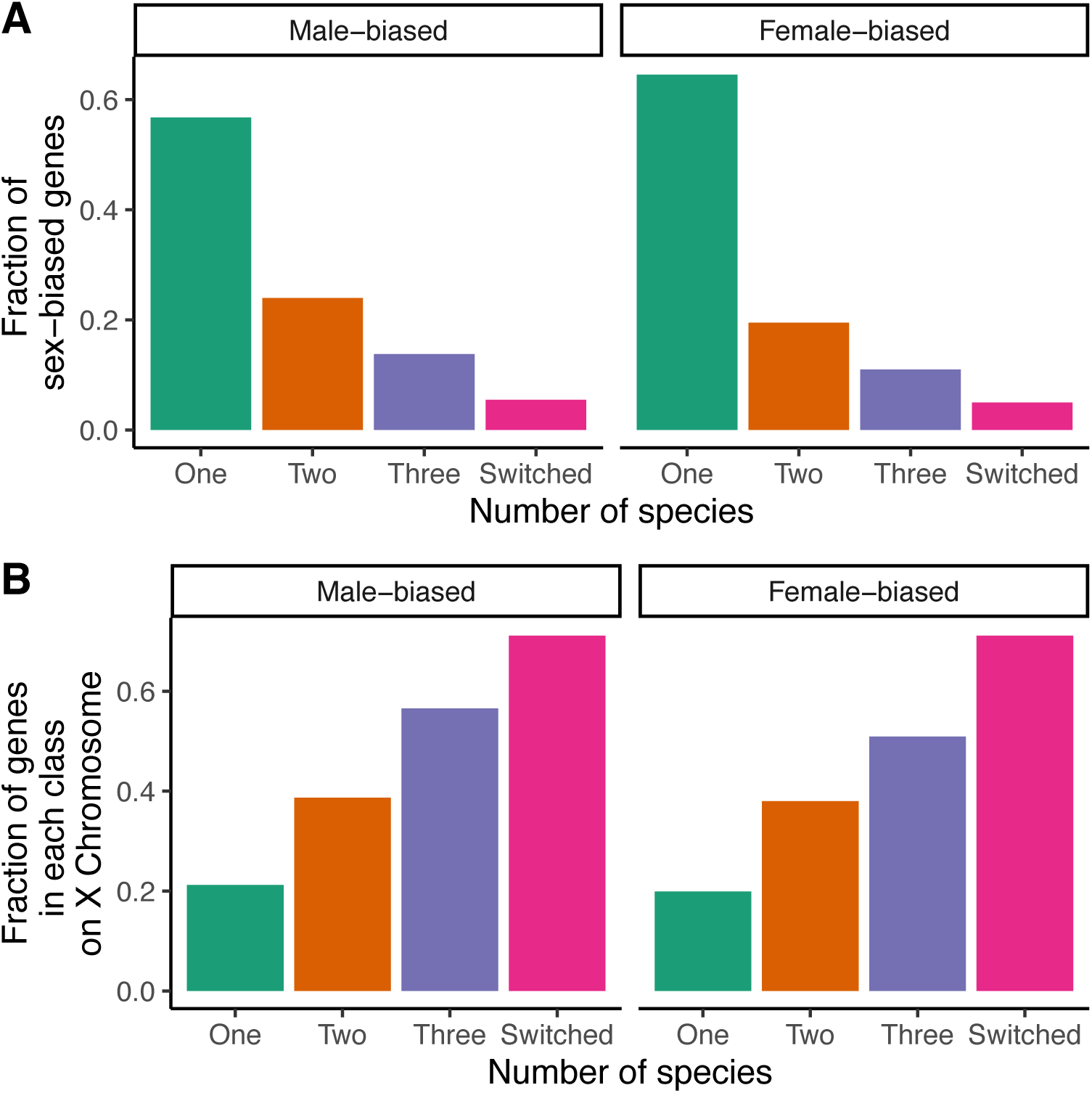
Expression turnover of sex-biased genes in the brain. A) The fraction of sex-biased genes that are sex-biased in 1, 2, or 3 species. Also shown is the fraction of sex-biased genes that have ‘switched’ sex-bias between species (i.e. genes which are male-biased in at least one species and female biased in at least one other species). B) The fraction of genes in each species number class (1, 2, 3, or switched) which are located on the X Chromosome. For any given gene the chromosomal location was taken to be the location of the *D. melanogaster* ortholog.

### Forces governing gene expression evolution in the brain are chromosome and sex-bias dependent

To infer the types of selection acting on gene expression levels in the *Drosophila* brain, we used an ANOVA framework, similar to the method described by Nuzhdin et al. (Nuzhdin et al., 2004). In particular, we took the mean sum of squares for species (*MS*_*species*_) as a quantification of interspecific variability in expression and the mean sum of squares for strain (*MS*_*strain*_) as a quantification of intraspecific variability in gene expression. We then calculated the ratio of *MS*_*species*_ to *MS*_*strain*_. Genes with higher ratios are presumably evolving under the influence of directional selection whereas genes with low ratios may be evolving under balancing selection. Genes with intermediate ratios require a closer examination of their levels of between- and within-species variability. Relatively high levels of both between- and within-species variability is indicative of relaxed selective constraint whereas genes with low interspecific and intraspecific variability are likely to be evolving under stabilizing selection (Nuzhdin et al. 2004).

Considering only 1:1:1 orthologs between the three species, we found that male-biased genes on the X Chromosome (defined as genes with male-bias in at least one species) were enriched for signatures of directional selection in both sexes, and female-biased genes (defined as genes with female-bias in at least one species) were enriched for signatures of directional selection in females (ratios and p-values see **Figure 4A, Supplementary Figure S4, Supplementary Table S4**). Female-biased autosomal genes showed lower than expected median values for *MS*_*species*_/*MS*_*strain*_, which is indicative of balancing selection or adaptive variation. To determine if there is a faster-X effect for gene expression evolution in the brain, we compared *MS*_*species*_/*MS*_*strain*_ between genes on the X Chromosome and the autosomes. We found that sex-biased genes showed evidence of faster-X evolution in both males and females (**Supplementary Table S5**). No effect was found for unbiased X-linked genes.

**Figure 4.**
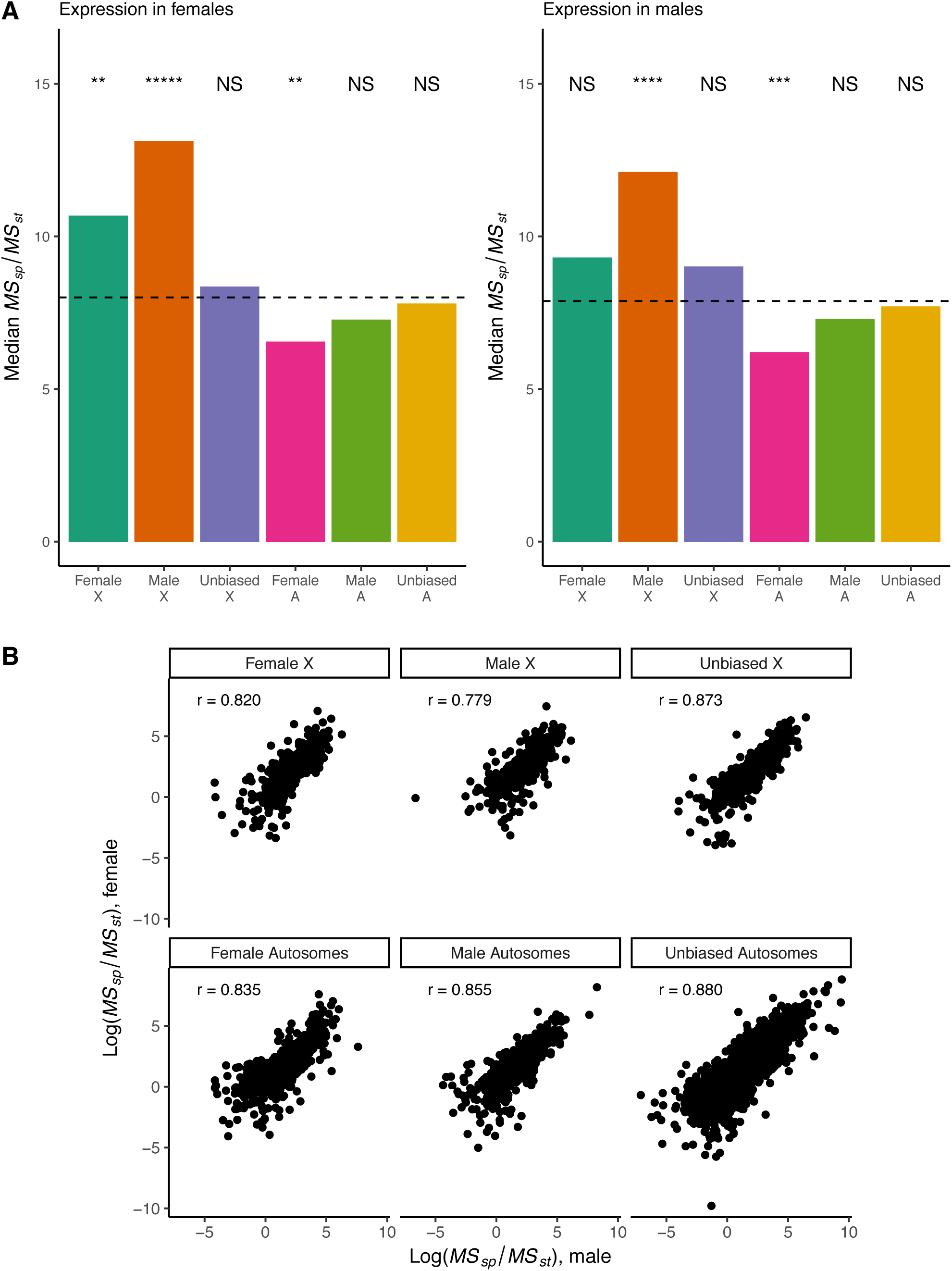
Ratio of interspecific to intraspecific variability in expression for genes in the *Drosophila* brain. A) Male-biased genes on the X Chromosome show significantly elevated variability ratios, consistent with directional selection. Similarly, female-biased genes on the X Chromosome show signatures of directional selection in females. Female-biased autosomal genes show significantly reduced ratios, consistent with balancing selection on gene expression. Significance of medians relative to the genome-wide median is shown. P-values determined using 1 million simulations per group and Benjamini-Hochberg correction. *****p-value < 1×10^−5^, ****p-value < 1×10^−4^, ***p-value < 1×10^−3^, **p-value < 1×10^−2^, *p-value < 5×10^−2^. A boxplot version of the data showing dispersion around the median is presented in Supplementary Figure 4. B) Ratios of interspecific to intraspecific variability in males (on x-axis) show large correlations (Spearman) with ratios in females (y-axis) irrespective of sex-bias or chromosomal location. Only 1:1:1 orthologs were considered. Genes were considered male- or female-biased if they exhibited that bias in at least 1 species at a FDR of 0.05. For any given gene the chromosomal location was taken to be the location of the *D. melanogaster* ortholog.

We then examined the marginal distributions of *MS*_*species*_ and *MS*_*strain*_ and noticed that male-biased autosomal genes showed lower than expected values for both statistics in both sexes, which is suggestive of stabilizing selection (**Supplementary Figure S5A, B**). To further investigate whether certain classes of genes were enriched for signatures of stabilizing selection, we examined the fold-enrichment of genes with lower-than genome-wide median values for both *MS*_*species*_ *and MS*_*strain*_. We found an enrichment of signatures of stabilizing selection in two gene classes: 1) male-biased autosomal genes in both sexes and 2) unbiased genes on the X Chromosome in males (**Supplementary Figure S6**). Interestingly, female-biased autosomal genes were depleted for signatures of stabilizing selection.

If the dosage compensation complex or other sex-specific *trans* acting factors were the main drivers of adaptive gene expression evolution for X-linked sex-biased genes, one would expect correlations between variability ratios in males and females to be fairly low. However, we observed a high correlation (Spearman’s r = 0.78 - 0.88) between variability ratios of sex-biased genes in males and in females (**Figure 4B**). This suggests that sex-specific *trans* acting factors are unlikely to be the main drivers of adaptive gene expression evolution for X-linked sex-biased genes.

To determine if the observed patterns of gene-expression evolution were associated with changes in sex-biased state, we compared genes with sex-bias in only one species with genes with conserved sex-bias in all three species (**Supplementary Figure S7**). With the exception of female-biased autosomal genes, most sex-biased genes showed similar patterns regardless of conservation.

Sex-biased genes with large *MS*_*species*_/*MS*_*strain*_ ratios (i.e. whose expression is likely evolving under directional selection) include the X-linked gene *cinnamon* (FBgn0000316, ratio_male = 86.15, ratio_female = 302.26). *Cinnamon*, which is involved in the biosynthesis of the molybdenum cofactor and inter-male aggression (Kamdar et al. 1994; Ramin et al. 2019), has changed sex-bias in its recent evolutionary history: it has female-biased expression in both *D. melanogaster* and *D. simulans* and male-biased expression in *D. yakuba*.

### Patterns of gene expression evolution are associated with the evolution of flanking sequences

To determine if DNA sequence evolution in gene-flanking regions was associated with the observed patterns of selection on gene expression, we examined the levels of nucleotide diversity in regions 20 kb upstream and downstream of sex-biased genes in *D. melanogaster.* Compared to unbiased genes and female-biased genes, we found a significant depression of nucleotide diversity (*π*) proximal to X-linked male-biased genes (**Figure 5A**). This indicates that these genomic regions may have recently been under the influence of positive selection, which is consistent with our observation of directional selection acting on these genes at the expression level. We then used the Hudson-Kreitman-Aguade-like (HKAl) statistic (Hudson et al. 1987; Begun et al. 2007) to look for further evidence that the flanking regions of sex-biased X Chromosome genes may be evolving under selection. Lower values of HKAl are indicative of positive selection. By considering the minimum HKAl value of genomic regions overlapping 20kb flanking regions of genes on the X Chromosome (excluding the genes themselves) we found that male-biased genes had significantly lower minimum HKAl values than both unbiased and female-biased X Chromosome genes (p = 0.001 and p = 0.004, respectively, two-sided Mann-Whitney *U* test) (**Supplementary Figure 8A**). Overall these results suggest that the observed enrichment of directional selection on X-linked male-biased genes is at least partly due to adaptive evolution in the flanking (putative regulatory) sequences of these genes.

**Figure 5.**
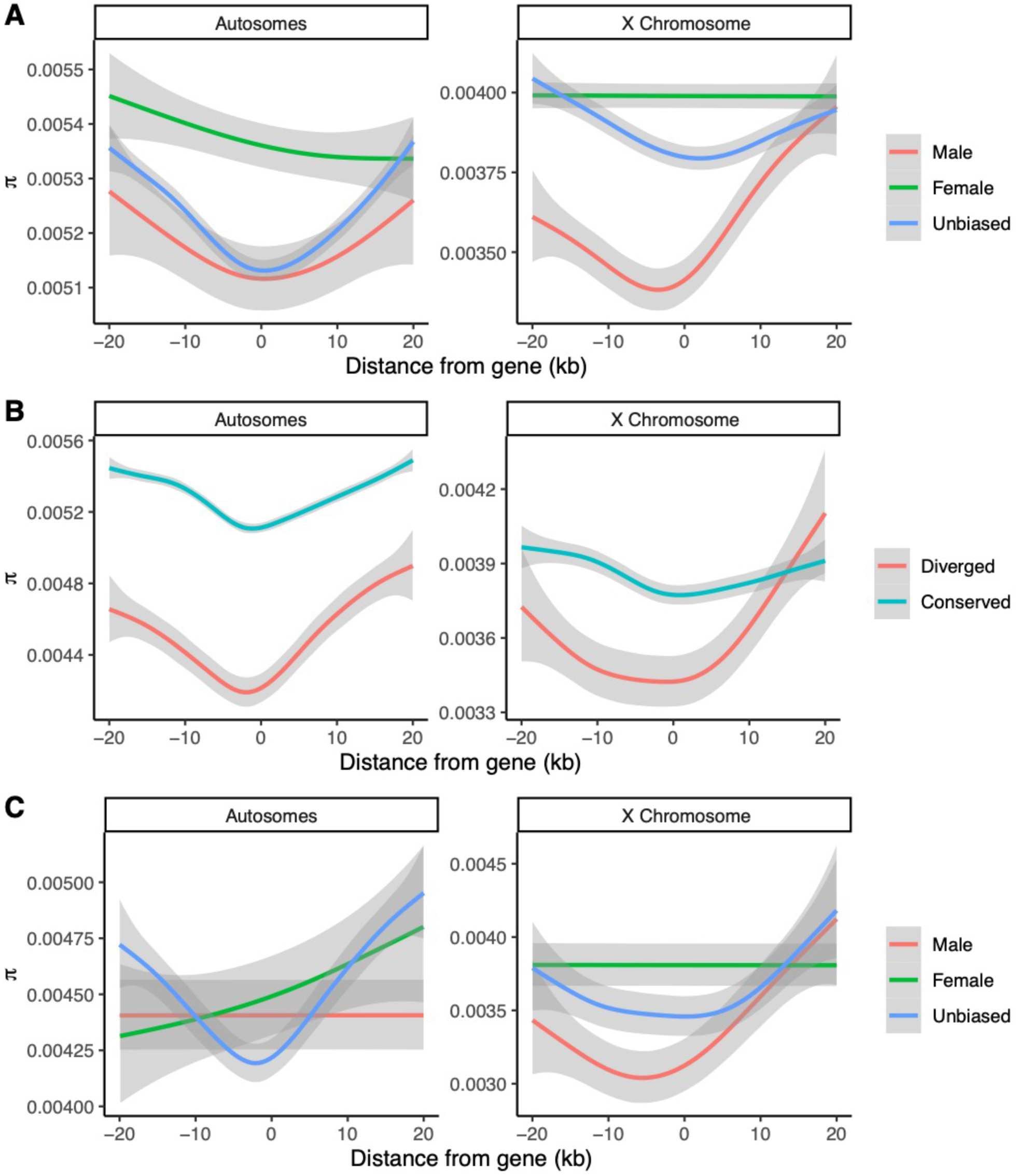
Nucleotide diversity in flanking regions of sex-biased genes. Nucleotide diversity (*π*) for various classes of genes on the X Chromosome and autosomes: A) sex-biased vs. unbiased genes, B) diverged vs. conserved genes, and C) diverged sex-biased vs. diverged unbiased genes. Diverged genes are defined as genes which are in the top 5% genome-wide in terms of *MS*_*species*_*/MS*_*strain*_ in either sex. Conserved genes are defined as genes which are not diverged in either sex. 95% confidence intervals are shown in gray. Sex-biased genes are defined in *D. melanogaster* at a FDR of 0.05.

We noticed that female-biased X Chromosome genes showed no clear adaptive signatures in their flanking regions despite showing signatures of adaptation in their expression (**Figure 4A and Figure 5)**. To explain this, we hypothesized that the expression evolution of male-biased X-linked genes is influenced by cis-regulatory changes to a greater extent than other classes of genes on the X Chromosome. When comparing the nucleotide diversities in the flanking regions of highly diverged sex-biased and unbiased genes (at the gene expression level, we defined diverged genes as genes that have ratios of *MS*_*species*_ to *MS*_*strain*_ in the top 5% genome-wide, conserved genes are not diverged in either sex), we observed that diverged male-biased genes on the X Chromosome had significantly reduced nucleotide diversity in their flanking regions when compared to other X-linked genes (**Figure 5C).** When comparing HKAl in the flanking regions of highly diverged sex-biased and unbiased genes we noticed a similar trend, however, the differences were not significant (**Supplementary Figure S8C**).

### Broad expression across tissues does not play a major role in constraining the directional expression evolution of genes in the *Drosophila* brain

To examine whether broad expression played a role in constraining expression evolution in the brain we calculated a tissue specificity statistic (Yanai et al. 2005) for genes expressed in males and females of both *D. melanogaster* and *D. yakuba* using published expression data from four tissues (abdomen, thorax, viscera, and gonads) (Yang et al. 2018), along with our brain data. A tissue specificity near 0 indicates broad expression whereas a tissue specificity near 1 indicates that expression is highly tissue specific. We found a very weak negative relationship between tissue specificity (in both sexes and species) and *MS*_*species*_*/MS*_*strain*_ in the brain, indicating that broad expression does not constrain directional gene expression evolution in the *Drosophila* brain (**Figure 6**).

**Figure 6.**
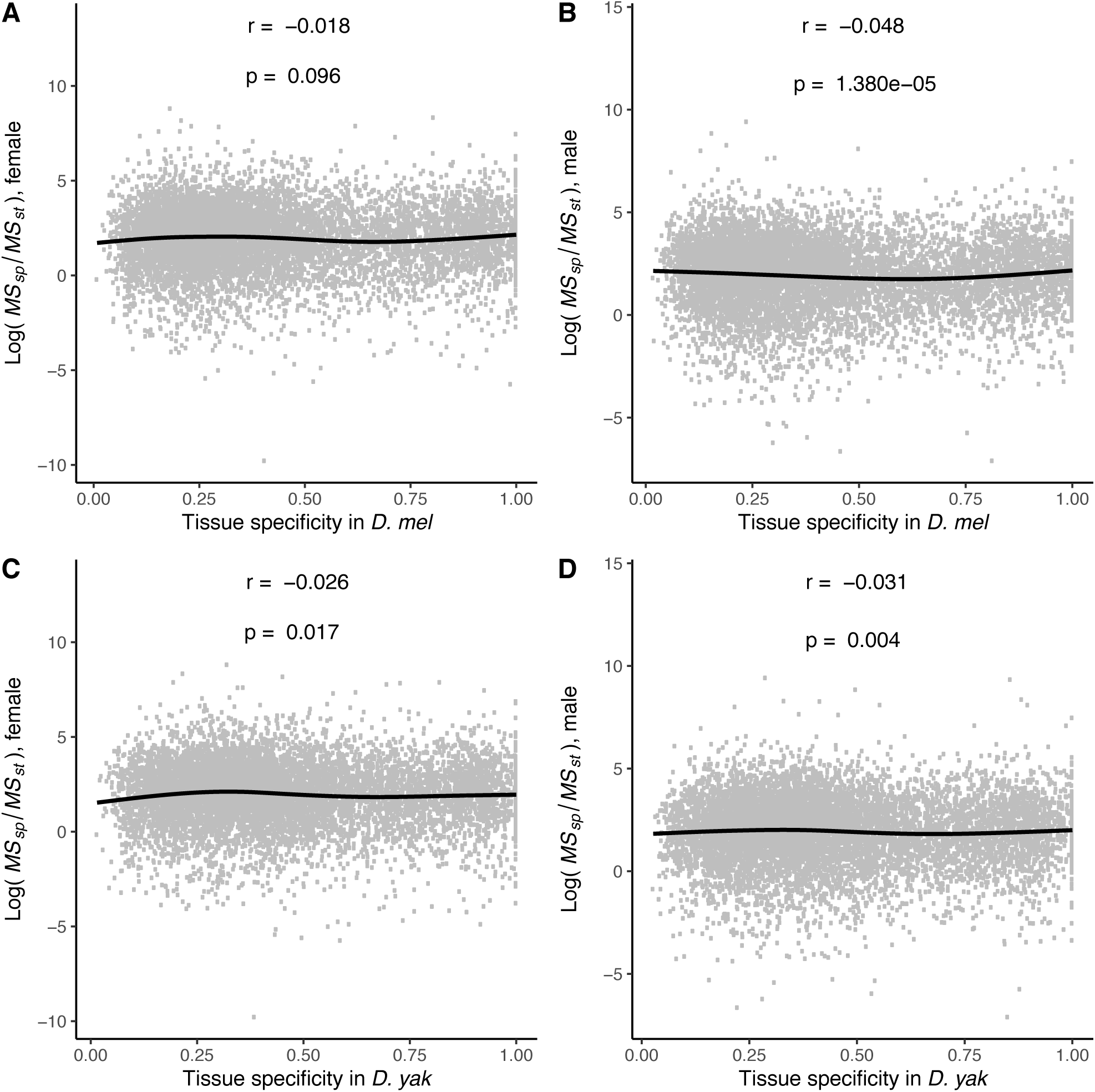
Expression breadth across tissues does not constrain gene expression evolution in the brain. There is a very weak but significant negative relationship between tissue specificity and *MS*_*species*_*/MS*_*strain*_ in the brain. The plots show the distribution of *MS*_*species*_*/MS*_*strain*_ for a range of tissue specificity in A) *D. melanogaster* females, B) *D. melanogaster* males, C) *D. yakuba* females, and D) *D. yakuba* males. A tissue specificity of 1 indicates expression in only one tissue and a low tissue specificity indicates broad expression. Correlations shown are Spearman correlations. Lines are generalized additive model smoothing curves. Sex-biased genes are defined in the same way as in Figure 4.

To determine if tissue specificity was associated with the observed patterns of directional expression evolution for sex-biased genes, we controlled for the effects of tissue specificity on *MS*_*species*_*/MS*_*strain*_ (**Supplementary Figure S9**). We found that all of the patterns we observed earlier (**Figure 4A, Supplementary Figure S4**) still held. Hence, it appears as though broad expression imposes little constraint on the directional evolution of gene expression in the brain and it cannot explain the signatures of directional selection observed for sex-biased genes on the X Chromosome.

Despite the observation that expression breadth does not constrain adaptive gene expression evolution, we wondered if it could be constraining gene expression variability. Therefore, we examined the association between tissue specificity and interspecific and intraspecific gene expression variability (*MS*_*species*_ and *MS*_*strain*_, respectively) (**Supplementary Figure S10**). We found that tissue specificity was positively correlated with both classes of gene expression variability in the brain. Broadly expressed genes tend to have low expression variability while tissue specific genes tend to have larger variability in expression.

To test whether expression breadth could explain the enrichment of signatures of stabilizing selection on the expression levels of autosomal male-biased and X-linked unbiased genes, we examined the tissue specificities of each gene class (**Supplementary Figure S11**). We observed that male-biased genes are generally broadly expressed and male-biased autosomal genes have particularly low tissue specificities. This suggests that expression breadth may partially explain the low inter- and intra-specific variability observed in male-biased autosomal genes. To examine this in greater depth, we controlled for the effects of tissue specificity on *MS*_*species*_ and *MS*_*strain*_ and ran our test for stabilizing selection (**Supplementary Figure S12**). The patterns we observed earlier (**Supplementary Figure S6**) were similar, but significance for male-biased autosomal genes was lost. This suggests that broad expression may be responsible for our observation of stabilizing selection on the expression levels of male-biased autosomal genes.

### The shared genome between the sexes constrains the expression evolution of sex-biased genes in the *Drosophila* brain

The shared genome has been shown to constrain the evolution of sexually dimorphic phenotypes in several species (Poissant et al. 2010; Griffin et al. 2013). An alternative to this is that sex-specific factors tend to dominate the expression evolution of these genes. To distinguish between these two scenarios, we examined changes in the expression levels of sex-biased genes relative to their ancestral state in both sexes. The ancestral expression level for a given gene was estimated by maximizing the likelihood of a Brownian motion evolutionary model on the three species phylogenetic tree. For both male-and female-biased genes in all three species, there was a strong correlation between the relative change in male expression and the relative change in female expression, indicating that male and female expression levels of sex-biased genes largely evolve in the same direction (**Supplementary Figure S13A, B**).

We then sought to examine the role of genetic constraint in shaping the expression of sex-biased genes on a gene-by-gene basis. To do so we looked at correlations between male and female expression levels for each sex-biased gene in *D. melanogaster*. Since males and females within a given strain possess nearly identical genomes, genes with largely positive correlations between male and female expression levels are genetically constrained. Regardless of sex-bias and chromosomal location, the distribution of correlations was heavily positively skewed, indicating that the shared genome imposes a significant constraint on the expression evolution of sex-biased genes (**Figure 7**). Given that we only have expression data from six *D. melanogaster* strains, it is likely that we consistently underestimated the correlations since it appears as though male-female correlations are generally strongly positive.

**Figure 7.**
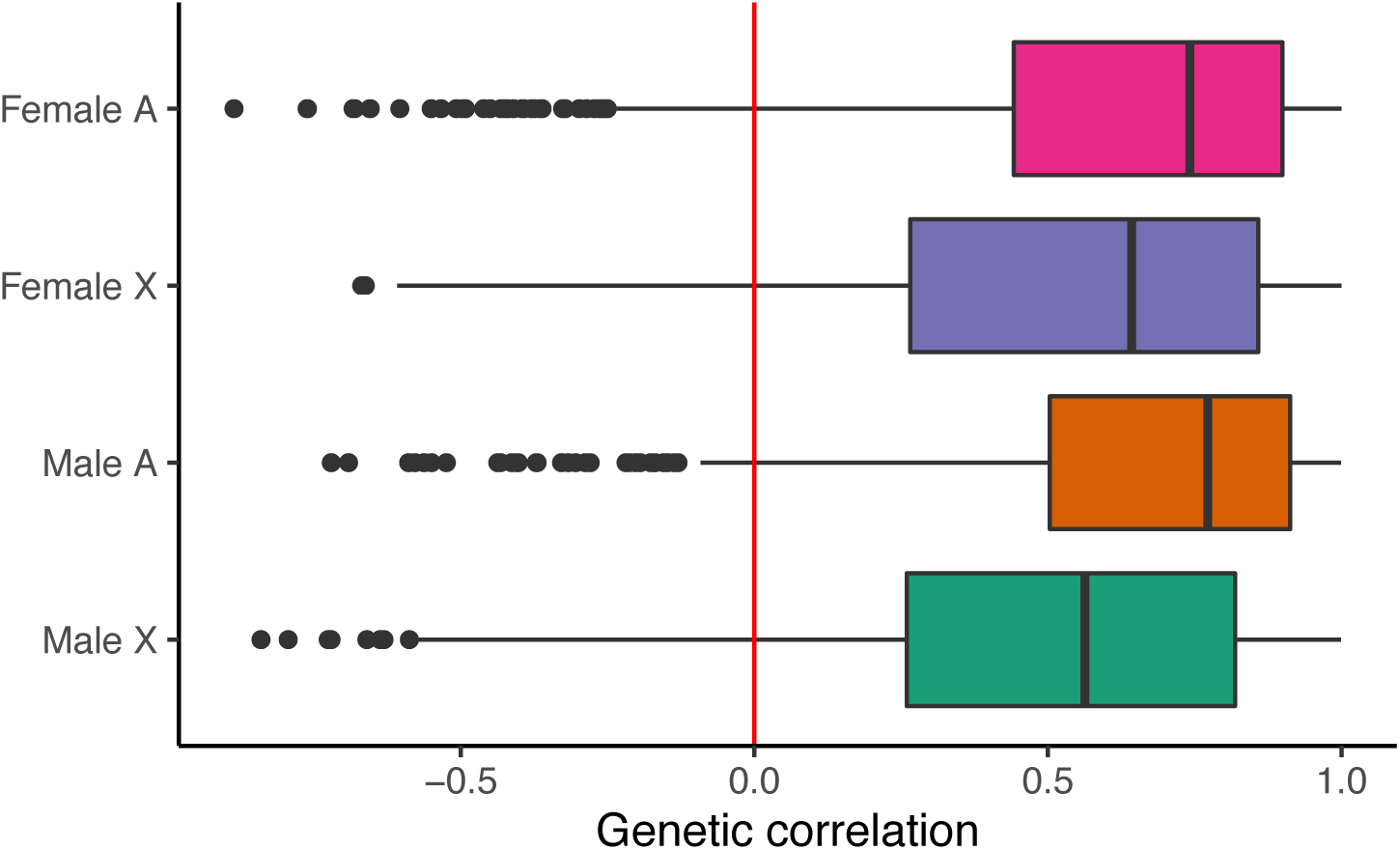
The shared genome constrains expression evolution of sex-biased genes in the *Drosophila* brain. Correlations between male and female expression levels for sex-biased genes in *D. melanogaster*. For each category (on the y-axis), a Pearson correlation coefficient (x-axis) was calculated for each gene using 6 male-female data points (one for each strain). The median correlations for each class of sex-biased gene are non-zero. One-sample Wilcoxon signed-rank test p-values < 2.2×10^−16^ for each class (female autosomal, female X, male autosomal, and male X. Sex-biased genes are defined in *D. melanogaster* at a FDR of 0.05.

### Evolution of sex-biased genes at the protein level in *D. melanogaster*

We then examined the forces governing the evolution of sex-biased genes at the protein level in *D. melanogaster*. First, we looked for evidence of selection on sex-biased genes by examining *d*_N_/*d*_S_ values generated from published data comparing *D. melanogaster* to *D. simulans* (Zhao and Begun 2017). *d*_N_/*d*_S_ is the ratio of nonsynonymous substitutions per nonsynonymous site to synonymous substitutions per synonymous site, with high ratios (>1) being indicative of positive selection and low ratios being indicative of purifying selection. We found that autosomal male-biased genes are significantly enriched for signatures of purifying selection (**Supplementary Figure S14**) This result, along with the relatively high expression of autosomal male-biased genes, relatively low sex-biased fold-change in all three species, low gene expression variability within and between species, broad expression, and enrichment of metabolism related GO terms suggests that autosomal male-biased genes have important, conserved functions in *Drosophila*.

To further understand whether positive selection has influenced the evolution of coding sequences of sex-biased genes, we examined published McDonald-Kreitman (MK) test data of annotated protein-coding genes in *D. melanogaster* (Langley et al. 2012). The MK test infers the presence of positive selection by comparing the numbers of fixed and polymorphic, synonymous and nonsynonymous substitutions (McDonald and Kreitman 1991). We found that sex-biased genes on the X Chromosome were significantly enriched, relative to the set of all genes, for genes with significant MK tests (**Supplementary Table S6A**). However, when comparing the numbers of genes with positive MK tests between sex-biased genes on the X Chromosome and all genes on the X Chromosome, we found that sex-biased genes showed a slight but non-significant enrichment (**Supplementary Table S6B)**.

## Discussion

We have carried out the first systematic study on the evolution of sex-biased gene expression in the *Drosophila* brain. The outcomes of our analysis in combination with past studies suggest that the evolutionary properties of sex-biased genes may be tissue specific. We found a significant enrichment of genes exhibiting male- and female-biased expression on the X Chromosome in three species of *Drosophila* (**Figure 2**). The X Chromosome harbored a significantly higher relative number of genes with conserved sex-biased expression in multiple species, suggesting that the X Chromosome acts as a ‘sink’ for sex-biased genes (**Figure 3**).

When examining the evolutionary forces acting on gene expression, we noticed that sex-biased X-linked genes were enriched for signatures of directional selection (**Figure 4A**). Moreover, when comparing X-linked genes with autosomal genes, we found evidence for a faster-X effect for the expression evolution of sex-biased genes (**Supplementary Table S5**). The assumptions underlying the most common explanation for faster-X evolution of gene expression are: the mutations largely influencing gene expression on the X Chromosome are X-linked, and the mutations are at least partly recessive with respect to expression or fitness (Meisel et al. 2012; Meisel and Connallon 2013). The signatures of adaptive evolution we observed in the flanking regions of X-linked genes – especially in male-biased genes – are consistent with the first assumption (**Figure 5**). Our observation of clearer signatures of adaptive evolution in genomic regions proximal to X-linked male-biased genes relative to other X-linked genes suggests that adaptive evolution in expression may be more likely to occur due to adaptation in *cis*-regulatory regions for male-biased genes than unbiased and especially female-biased genes (**Figure 5C**). This is consistent with previous findings suggesting that adaptive evolution of *cis*-regulatory regions for male-biased X-linked genes, in particular, could be partly responsible for a faster-X effect in expression for those genes (Coolon et al. 2015). Additionally, it has been observed that *trans* factors responsible for the expression of X-linked genes tend to disproportionally reside on the X Chromosome (Coolon et al. 2015). Therefore, it is possible that evolution in *trans*-regulatory regions is partly responsible for adaptive expression evolution in sex-biased genes. Given that we observed a faster-X effect for expression in female-biased genes but not in unbiased genes may indicate that *trans* factors regulating the expression of X-linked female-biased genes may be more likely to reside on the X Chromosome than *trans* factors regulating unbiased X-linked genes.

We examined the role that broad expression and shared genomic constraints impose on the expression evolution of sex-biased genes and found that broad expression across tissues generally does not constrain the directional evolution of gene expression in the *Drosophila* brain (**Figure 6**). However, we found that expression breadth does appear to constrain expression variability, both between and within species. The fact that expression breadth does not seem to inhibit directional gene expression evolution suggests that expression evolution in the brain might in part be attributable to selection acting on expression in other tissues which in turn ‘drag along’ expression levels in the brain and vice versa.

In accordance with findings made in other systems, we found that the shared genome between the sexes significantly constrains the expression evolution of sex-biased genes. This indicates that selection acting upon the expression level of a certain sex-biased gene in one sex will likely ‘drag along’ the expression of that gene in the other sex. It has previously been shown that the chromatin structures of the X Chromosome in male and female *Drosophila* are similar, despite the fact that dosage compensation only occurs in males, and this might be because the need for dosage compensation in males imposes a constraint on the chromatin structure in females (Zhang and Oliver 2010). This suggests that dragging between sexes can even occur at the epigenetic level. Of course, dragging between sexes and tissues is only likely to occur if the effects of dragging are beneficial, neutral, or only slightly deleterious in the ‘dragged’ tissue or sex. We speculate that gene expression dragging between sexes and tissues is a pervasive phenomenon that may be responsible for various observed instances of phenotypic evolution. If this is indeed the case, it will significantly alter our understanding of gene expression evolution, and needs to be investigated in depth in the future.

The genes which we identified as rapidly evolving at the expression level may be thought of as candidate genes involved in the evolution of species-specific neural circuits and behaviors. Therefore, future research should systematically investigate the role that these genes play in rapidly evolving neural circuits and behaviors in *Drosophila*, both sexually dimorphic and otherwise (Seeholzer et al. 2018). Additionally, since mating alters gene expression patterns in *Drosophila* heads (Ellis and Carney 2010; Dalton et al. 2010), it would be interesting to see how the results of a study of sex-biased expression in unmated or young flies would compare to our results. Ultimately, the long-term goal is to reveal how sex-biased gene expression in somatic tissues contributes to reproduction and fitness in a species.

## Methods

### Fly lines and tissue collection

We collected data from inbred strains of three *Drosophila* species: the *D. melanogaster* strains come from the DGRP panel which are wild caught flies from Raleigh, NC, USA: RAL304, RAL307, RAL357, RAL360, RAL 399, RAL517 (MacKay et al. 2012); *D. simulans* strains were FAIR3 (Zhao et al. 2015) and *w*^*501*^ (Begun et al. 2007); *D. yakuba* strains were Tai18E2 (Clark et al. 2007) and CY07, which is a Cameroonian isofemale line (Bachtrog et al. 2006). All lines were maintained on standard media at 24°C under a 12:12 light/dark cycle.

Prior to the dissections, all fly lines were expanded and maintained at 24°C under an undisturbed 12:12 light/dark cycle in low density vials with fly food. All individuals were mated and precisely 4-days of age. Flies were very briefly anesthetized with CO_2_, and heads were collected with a clean razor blade, then were transferred to a glass dissection plate with PBS for brain collection. Dissections were performed within a 3-hour window always at the same circadian zeitgeber time (ZT1-ZT4; in this experiment the light was turned on at 8 am, and the dissections were performed between 9 am and 12 pm within a three-hour interval). Male and female brains of the same line were dissected in parallel the same day at the same time. Dissected brains were immediately transferred into a low retention Eppendorf tube containing 100 μL Trizol (Invitrogen) and kept frozen at −20°C until RNA extraction. We collected 3 biological replicates for each genotype by sex combination, and each replicate comprised of a pool of exactly 15 brains.

### RNA extractions and library preparation

All RNA extractions were performed according to Trizol manufacturer protocol and immediately followed by a DNase treatment using the TURBO DNase from Invitrogen. RNA quality was assessed by a Bioanalyzer run of an Agilent Eukaryote Total RNA Pico chip while RNA quantity was measured with a Nanodrop One (ABI). About 50 ng total RNA was used for library preparation. Libraries were fragmented and enriched for mRNA using NEBNext Poly(A) Magnetic Isolation Module (NEB #E7490) and prepared using NEBNext Ultra II Directional RNA Library Prep Kit (NEB #E7765) and dual indexing from the NEBNext Multiplex Oligos kit (NEB #7600) following manufacturer protocol including beads size selection for 200 bp inserts. Library quality was first assessed on Agilent D1000 ScreenTapes for Tapestation and then by Qubit and Agilent Bioanalyzer. Finally, 150 bp paired-ends libraries were sequenced on 5 lanes of an Illumina Nextseq500 platform (**Supplementary Table S7**).

### Read mapping and identification of sex-biased genes

Adaptors from RNA-seq reads were trimmed using Trimmomatic (Bolger et al. 2014). In addition, low-quality bases were trimmed using Trimmomatic settings LEADING:1 TRAILING: 1 SLIDINGWINDOW:20:25 and MINLEN:36. HISAT2 (Sirén et al. 2014) was then used to align reads to their respective reference genomes: dmel_r6.15 for *D. melanogaster*, dsim_r2.02 for *D. simulans*, and dyak_r1.05 for *D. yakuba.* Gene and transcript expression levels (TPM: Transcripts per million) were then quantified using StringTie and read counts were extracted using a script, prepDE.py, provided by StringTie (Pertea et al. 2015). We performed quality control on the data obtained from the 60 libraries: three of them (one replicate of male FAIR3, male R517, and female R360) were excluded from further analysis because of low correlation with other replicates or deviation from other samples in the PCA plot.

The read counts were then analyzed using DESeq2 (Love et al. 2014) to identify sex-biased genes in each species. Genes with fewer than 10 read counts (summed across samples within a species) were excluded from the analysis. The DESeq2 model used to determine differential expression between the sexes was: counts ∼ sex + strain. Genes with statistically significant ‘sex’ terms (FDR <0.05) were considered to be sex-biased in the given species. In all future analysis where we compared sex-biased genes to unbiased genes, we considered genes to be unbiased if DESeq2 assigned an insignificant p-value to the sex term for a gene. Genes with sufficiently low read counts such that the p-value was ‘NA’ were not considered.

To directly compare gene expression levels between samples from different species we used TMM (trimmed mean M-values) normalized TPM values. TMM normalization was implemented with the ‘R’ package edgeR (Robinson et al. 2010).

### Chromosomal distributions of sex-biased genes

To determine significance for the enrichment of sex-biased genes on the X Chromosome we used random sampling. For each simulation the total number of male- and female-biased genes was taken to be the same as the number observed at a FDR of 0.05. That number of genes was then randomly sampled from a list of all genes with finite p-values (from DESeq2), and the number of sampled genes on each chromosome arm was recorded. Simulations were repeated 100,000 times for each bias-species combination. The p-values were calculated as 2 times the fraction of simulations which resulted in a more extreme number of sex-biased genes on a given chromosome arm.

### Expression turnover of sex-biased genes

To get a conservative estimate for the expression turnover of sex-biased genes, equal numbers of genes were taken to be sex-biased in each of the three species. This was done by increasing the FDR in *D. simulans* and *D. yakuba* so that the numbers of sex-biased genes in each species was the same as the number found in *D. melanogaster* at a FDR of 0.05. Within these three sets we subsequently only considered 1:1:1 orthologs identified using a list obtained from FlyBase (Thurmond et al. 2019). Genes which are sex-biased in 1, 2, or all 3 species were identified, as well as genes which ‘switched’ sex-bias in their recent evolutionary history. ‘Switched’ genes are genes which are male-biased in at least one species and female-biased in at least one other species.

### Gene expression evolution

One-to-one-to-one orthologs between the three species were identified using a list obtained from FlyBase (Thurmond et al. 2019). TMM normalization of TPM values was then applied across all samples and replicates simultaneously for orthologous genes. Only genes with expression higher than 1 TPM in at least one of the three species were considered for further analysis. The resulting expression values were then log_2_ transformed and Pareto scaled. For every single gene we fit a model expression ∼ species + strain. Strain was taken to be a random effect and was nested within species. The model was fit using the MIXED procedure in SAS. The model was fit using the type III sum of squares. We then took the ratio of mean sum of squares (*MS*) for the species term as a quantification of interspecific variability and *MS* for the strain term as a quantification of intraspecific variability. A high value for the ratio *MS*_*species*_*/MS*_*strain*_ was taken as indicative of directional selection on expression while low values of the ratio were taken as indicative of balancing selection. To compare ratios, interspecific variability and intraspecific variability of various gene classes to genome-wide medians were used for simulations. We randomly sampled gene sets of identical size to the gene set of interest and computed medians for the statistic of interest. Simulations were run 1 million times, for each set of interest, and p-values were defined as twice the proportion of simulations where the median value for the statistic of interested was more extreme than the observed median. Genes were considered ‘male-biased’ or ‘female-biased’ if they were male-biased or female-biased in at least one of the species. Genes which were male-biased in at least one species and female-biased in at least one other species were removed from the analysis.

### Calculation of tissue specificity

RNA-seq data from the thorax, abdomen, gonads, and viscera of *D. melanogaster* and *D. yakuba* were obtained from Yang et al. (Yang et al. 2018). Reads were aligned and expression was quantified in the same manner as for our brain data (except adaptors were not trimmed before alignment). Within each species, sex, and tissue (including our brain data) we computed median TPM values which were then log_2_ transformed. Tissue specificity (*τ*) was then calculated in each sex and species using the formula 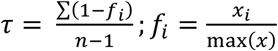, where *x*_*i*_ is the expression of gene *x* in tissue *i* (Yanai et al. 2005). The sum and maximum are taken over all tissues with *n* being the total number of tissues.

### Controlling for tissue specificity

To control for the effects of tissue specificity from *MS*_*species*_*/MS*_*strain*_, *MS*_*species*_ and *MS*_*strain*_ we used a generalized additive model fit to the relationship between log(‘statistic’) and tissue specificity (*τ*), where ‘statistic’ is the statistic of interest. To do so, we used this we used the function ‘gam’, from the ‘R’ library mgcv, with formula y ∼ s(x, bs=“cs”). The residuals were then calculated and plotted.

### Direction of gene expression evolution and the shared genome constraint

To infer ancestral expression levels for a given gene, we maximized the likelihood of a Brownian motion model on the three species phylogenetic tree. Since our results provide evidence that sex-biased genes are enriched for signatures of selection at the expression level, we only used a Brownian model as an approximation. In particular we used TMM normalized and log_2_ transformed expression levels (across all samples in all species) in combination with the anc.ML function from the ‘R’ library phytools to infer the ancestral expression levels. This was done for sex-biased genes with 1:1:1 orthologs in all three species. Additionally, we required that all genes considered had a median expression TPM>1 in at least one species in each sex. The ancestral state was estimated independently for males and females. For a given gene, the change in expression relative to its ancestral state was defined as (current – ancestral)/abs(ancestral), where ‘current’ is the log_2_(TPM + 1) of the extant expression and ‘ancestral’ is the log_2_(TPM + 1) of the inferred ancestral expression. Pearson’s correlations between male and female changes relative to ancestral state were then calculated.

To investigate the role of genetic constraint on a gene-by-gene basis, we first TMM normalized the *D. melanogaster* TPM data. Then for each sex, strain, and gene we calculated median expression, leading to 6 male-female pairs for each gene. Using these 6 data points (per gene) we calculated Pearson’s correlations for each sex-biased gene in *D. melanogaster*.

### Analysis of nucleotide diversity in gene flanking regions

SNP data for *D. melanogaster* was obtained from DGRP freeze2 (MacKay et al. 2012). Nucleotide diversity (*π*) within all DGRP strains was then calculated genome-wide in 500bp windows using VCFtools (Danecek et al. 2011). Genes were considered ‘diverged’ at the expression level if they had a ratio of *MS*_*species*_*/MS*_*strain*_ in the highest 5% genome-wide in either sex. For each gene in a given class (i.e. male-biased X Chromosome, unbiased autosomes, etc.), *π* windows overlapping +/- 20kb flanking regions were considered. The *π* of the gene is represented as a single point at 0. All of the data (distance from gene for each *π* window and the value of *π* within each window) from each gene in a given class was aggregated and generalized additive model smoothing was performed. To smooth the data we used the function ‘gam’, from the ‘R’ library mgcv, with formula y ∼ s(x, bs=“cs”), distribution family ‘Gamma’, and link function ‘log’. The resulting curves were plotted with the function ‘geom_smooth’ from the ‘R’ library ‘ggplot2’.

### Protein evolution analysis

Protein coding genes and orthologous information were downloaded from FlyBase (Thurmond et al. 2019). One-to-one orthologous genes in *D. melanogaster* and *D. simulans* were aligned using Genewise to remove insertions and deletions, then re-aligned using PRANK with the – codon option (Zhao and Begun 2017). We then used a KaKs Calculator (Zhang et al. 2006) to calculate *d*_N_ and *d*_S_ for aligned gene pairs between each of the two species using YN model (Yang and Nielsen 2000). The *D. melanogaster* McDonald-Kreitman test results including the Raleigh and Malawi samples was reported in Langley et al. (Langley et al. 2012). We estimated the proportion of adaptive amino acid fixations (α) of each gene according to the Smith and Eyre-Walker (Smith and Eyre-Walker 2002) and the Direction of Selection (DoS) index according to Stoletzki and Eyre-Walker (Stoletzki and Eyre-Walker 2011).

### Data and materials availability

The RNA-seq data generated in this study have been submitted to NCBI’s BioProject database (https://www.ncbi.nlm.nih.gov/bioproject/) under accession number PRJNA553234.

## Acknowledgments

We are extremely grateful to Roger Vaughan and Caroline Jiang for their help with the ANOVA analysis. We thank David Begun for sharing the *Drosophila* strains. We thank Grace Y.C. Lee and Jen Perry for critical reading of the manuscript, and members of the Zhao laboratory for helpful discussions during the work.

## Author contributions

L.Z., N.S., and S.K. conceived the project. L.Z. and N.S. designed the experiments. N.S. and S.D. generated the data. S.K. performed data analysis with the input from L.Z. and N.S.. S.K. and L.Z. wrote the manuscript with the input from all authors.

---

**Supplementary Figure S1.**
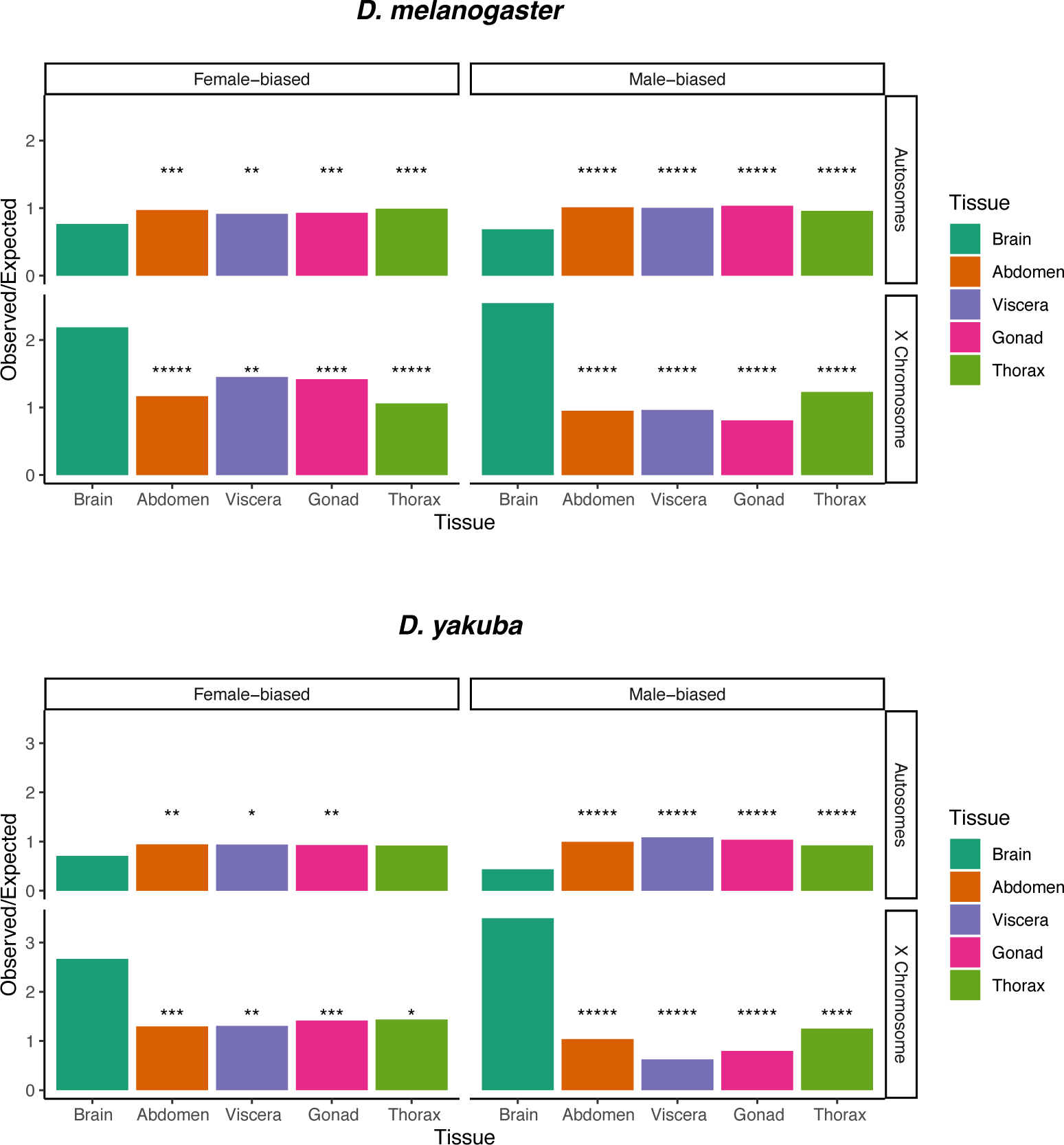
Fold enrichment of sex-biased genes on the X Chromosome and autosomes – a comparison between various tissues and the brain. A comparison of the number of observed vs. expected sex-biased genes between the brain and other tissues. Sex-biased genes in each tissue were found at a FDR of 0.05. P-values determined using a Chi-squared test followed by Benjamini-Hochberg correction *****p-value < 1e-5, ****p-value < 1e-4, ***p-value < 1e-3, **p-value < 1e-2, *p-value < 5e-2. Sex-biased genes are defined in each species at a FDR of 0.05.

**Supplementary Figure S2.**
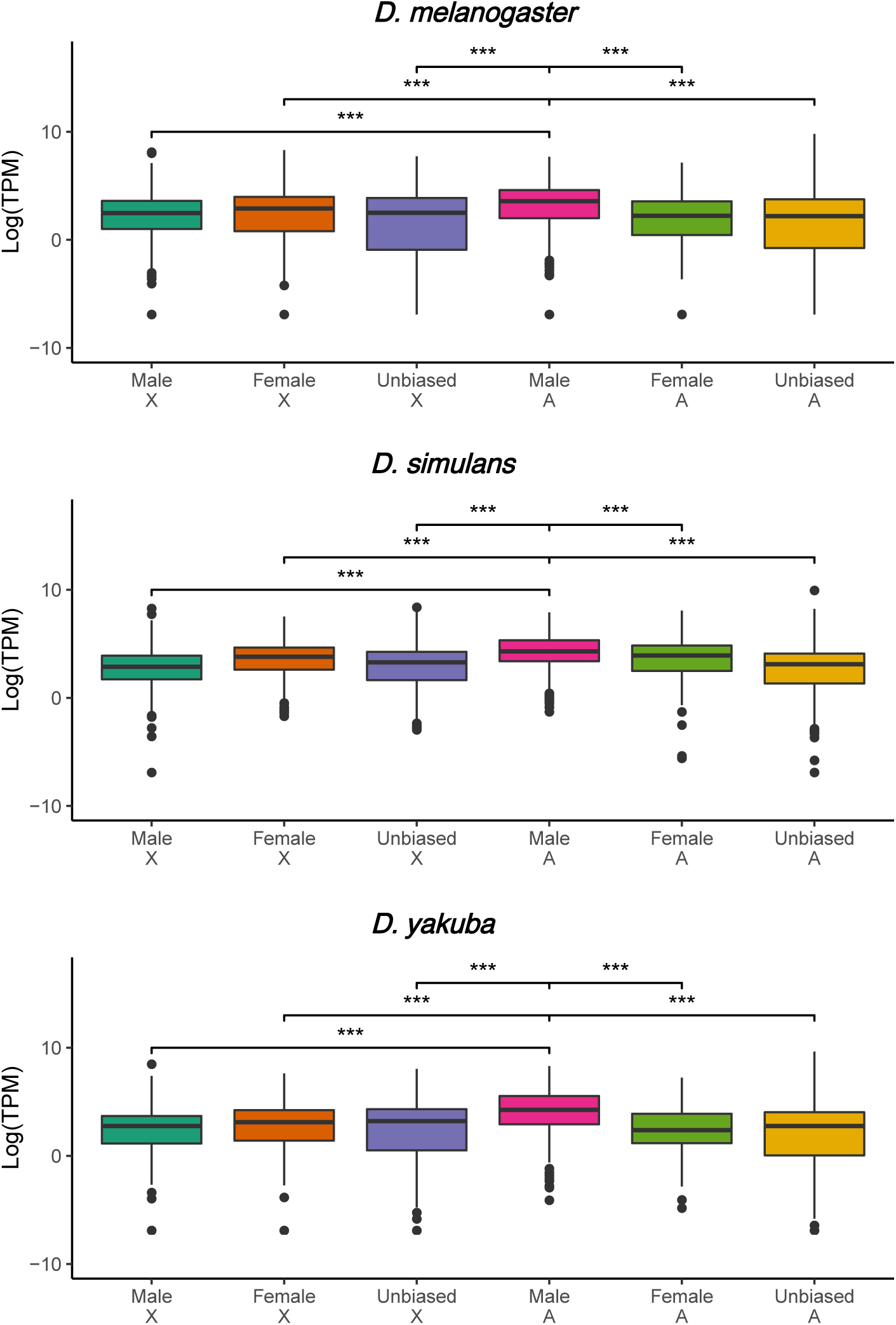
Expression levels of sex-biased genes in the *Drosophila* brain. Male-biased autosomal genes have higher expression levels than other classes of genes. ***p-value < 0.001. Two-sided Mann-Whitney *U* test. Expression levels were taken to be the median across all samples (both male and female) in each species. Sex-biased genes are defined in each species at a FDR of 0.05.

**Supplementary Figure S3.**
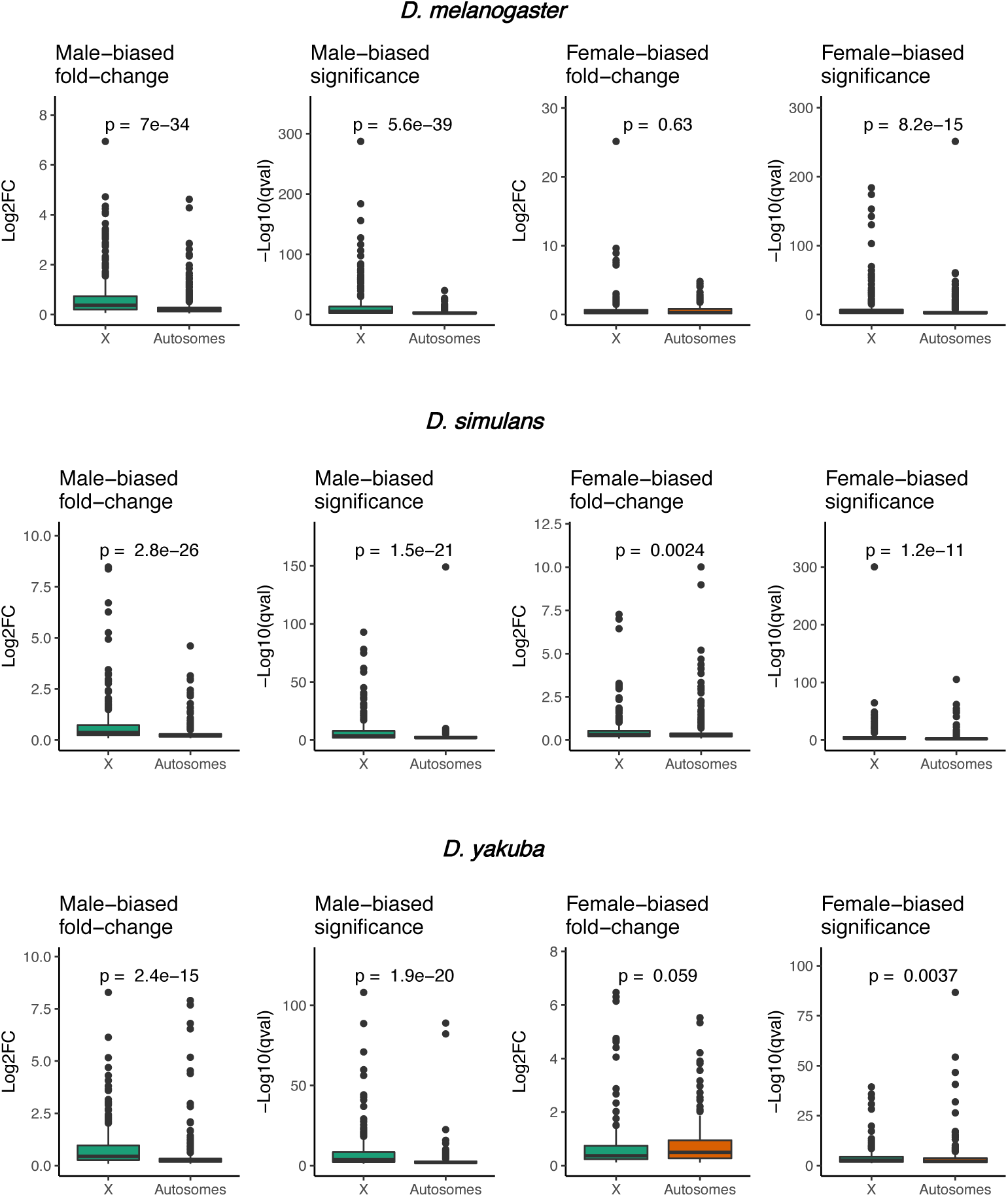
Log_2_ fold-change and −Log_10_ q-value for genes with sex-biased expression in the *Drosophila* brain. Male-biased genes on the X chromosome display both significantly larger fold-change and significance than male-biased autosomal genes. However, female-biased genes on the X Chromosome generally do not display a larger fold change than autosomal genes, but they are more significant. This suggests that female-biased autosomal genes may have relatively noisy expression. P-values were calculated using two-sided Mann-Whitney *U* test. Sex-biased genes are defined in each species at a FDR of 0.05.

**Supplementary Figure S4.**
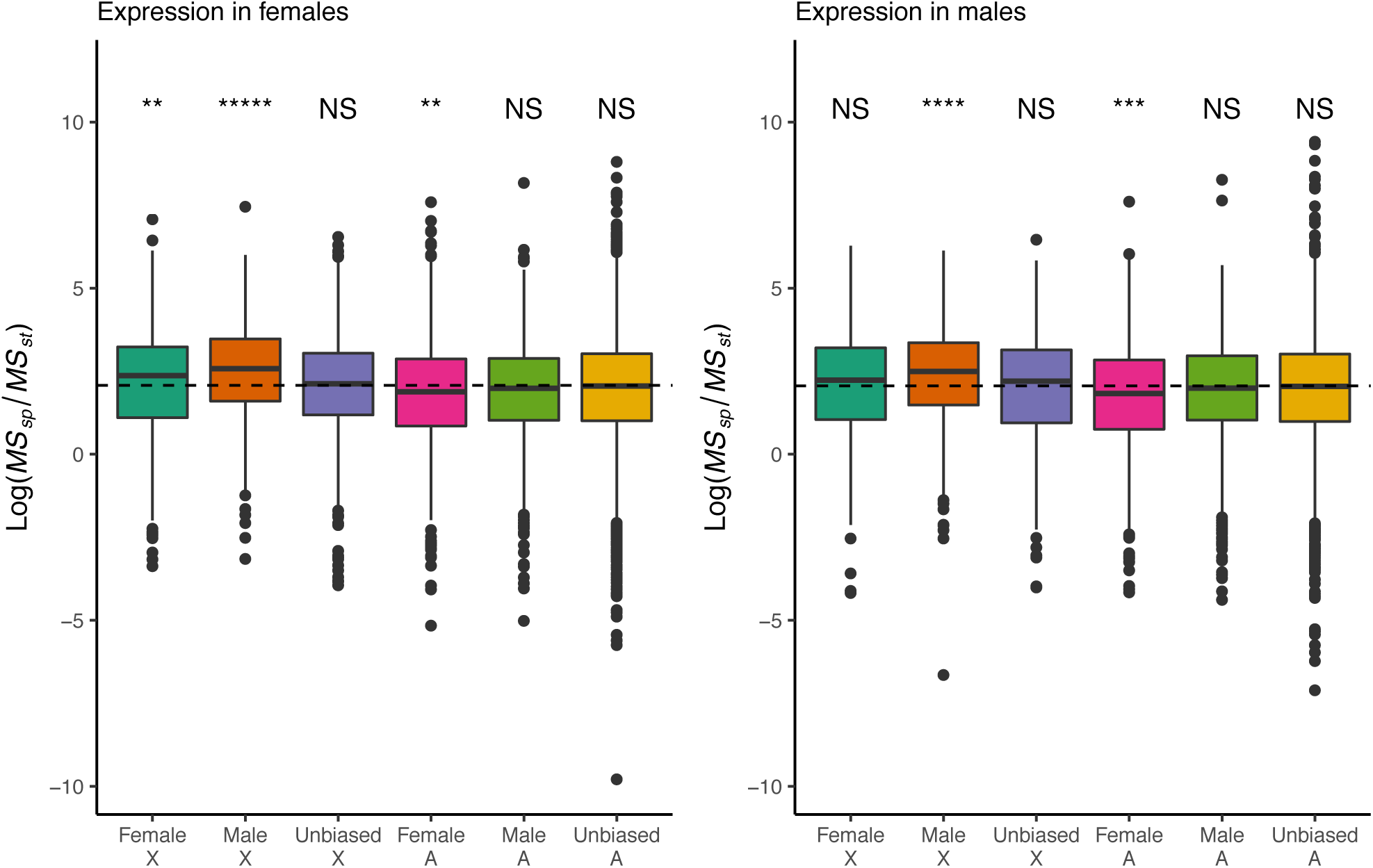
Ratio of interspecific to intraspecific variability in expression for genes in the *Drosophila* brain. Same data shown as in Figure 4A but with values log-transformed and data shown in boxplot form.

**Supplementary Figure S5.**
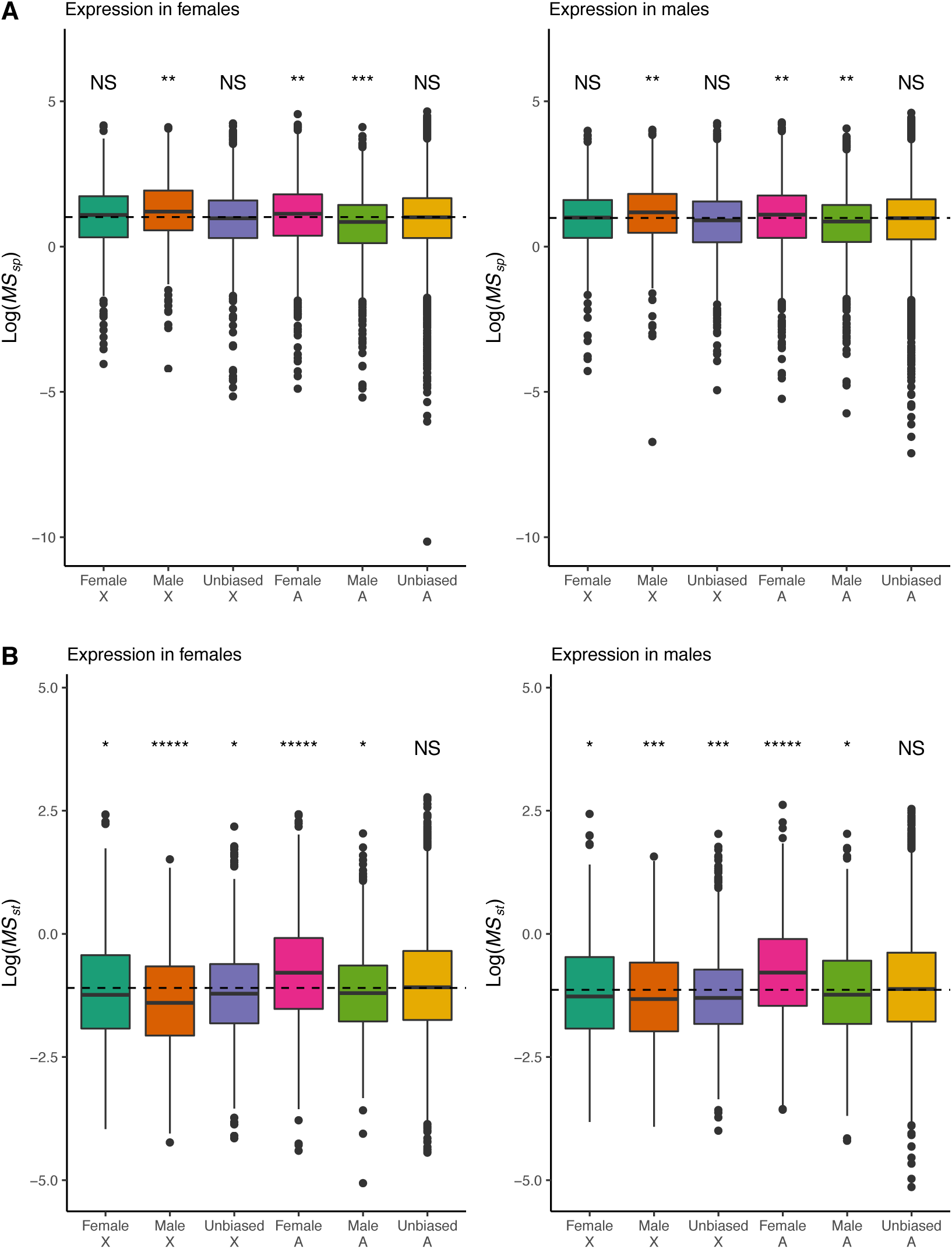
Marginal distributions of interspecific and intraspecific variability for genes in the *Drosophila* brain. A) Interspecific variability in gene expression, log(*MS*_*species*_). Female-biased autosomal and male-biased X Chromosome genes have relatively high interspecific variability in expression in both sexes. Male-biased autosomal genes have relatively low interspecific variability in expression in both sexes. B) Intraspecific variability in gene expression, log(*MS*_*strain*_). Male-biased autosomal genes have relatively low intraspecific variability in expression in both sexes. Female-biased autosomal genes have relatively high intraspecific variability in expression in both sexes. Significance of medians relative to the genome-wide median is shown. P-values determined using 1 million simulations per gene-class and Benjamini-Hochberg correction. *****p-value < 1e-5, ****p-value < 1e-4, ***p-value < 1e-3, **p-value < 1e-2, *p-value < 5e-2. Sex-biased genes defined in the same way as in Figure 4.

**Supplementary Figure S6.**
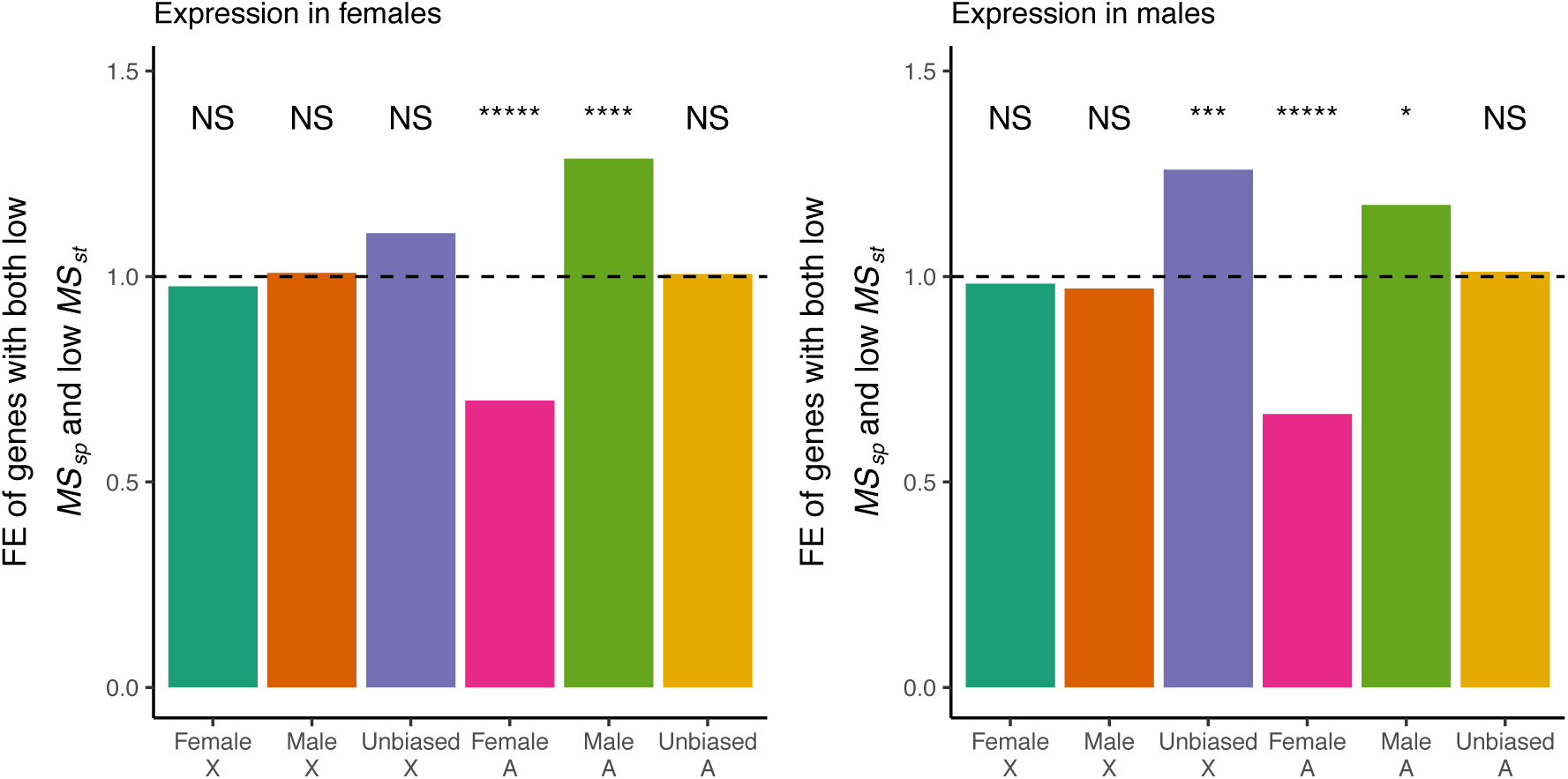
Fold enrichment of genes with both low intraspecific and low interspecific variabilities. Genes with low interspecific and intraspecific variabilities were defined as genes with MS_species_ and MS_strain_ both below the genome-wide medians. P-values were determined using a hypergeometric test followed by Benjamini-Hochberg correction. *****p-value < 1e-5, ****p-value < 1e-4, ***p-value < 1e-3, **p-value < 1e-2, *p-value < 5e-2. Sex-biased genes defined in the same way as in Figure 4.

**Supplementary Figure S7.**
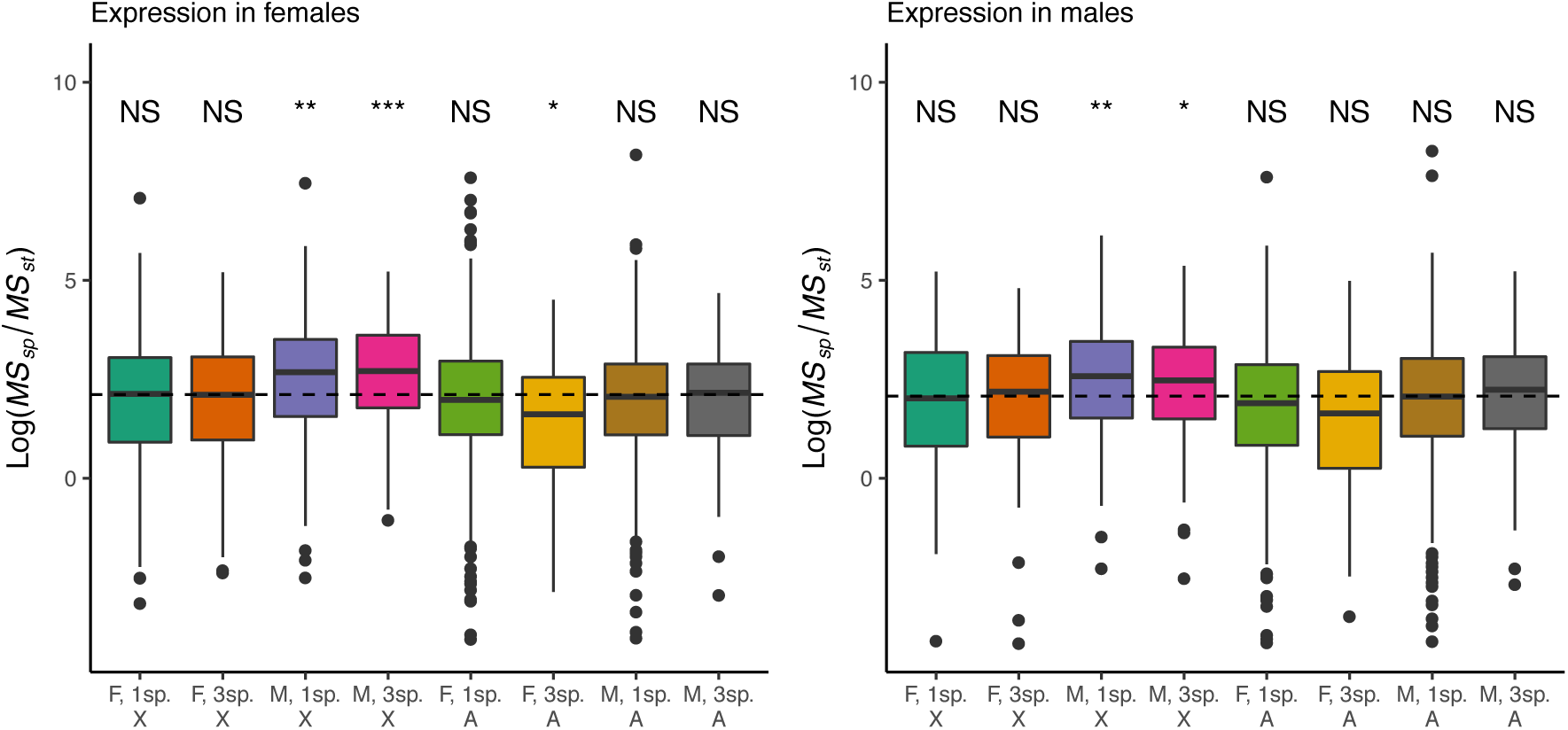
Ratio of interspecific to intraspecific variability in expression for genes with conserved sex-bias in all three species and genes with sex-bias in only one species. The ratio of interspecific to intraspecific variability is similar for genes with conserved sex-bias in all three species and genes with sex-bias in only one species. Significance of medians relative to the genome-wide median is shown. P-values determined using 1 million simulations per group and Benjamini-Hochberg correction. *****p-value < 1e-5, ****p-value < 1e-4, ***p-value < 1e-3, **p-value < 1e-2, *p-value < 5e-2. Sex-biased genes defined in the same way as in Figure 4.

**Supplementary Figure S8.**
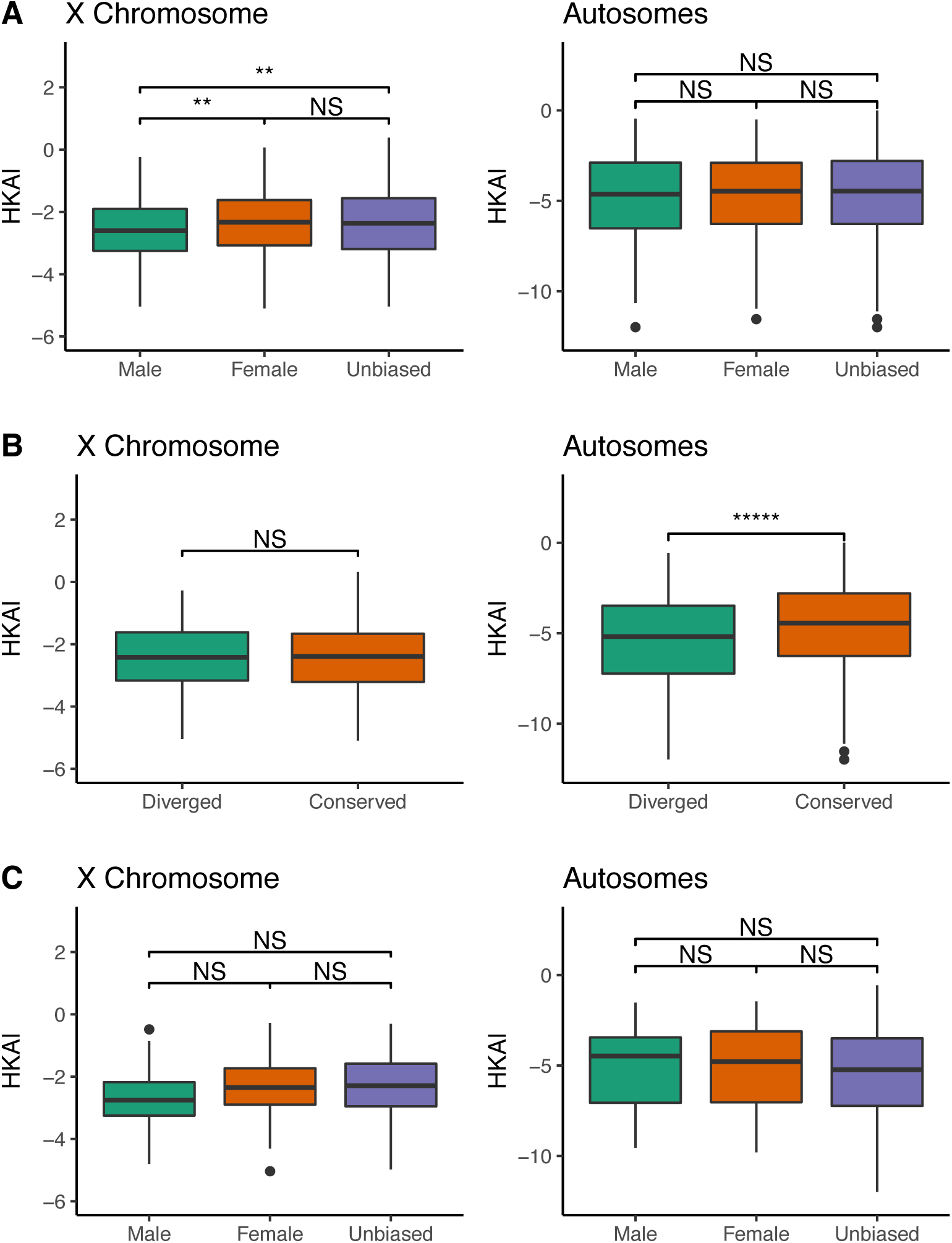
The minimum Hudson-Kreitman-Aguadé-like (HKAl) statistic value within 20kb of each gene. The statistic values only include flanking regions but not genes themselves. A) Comparison of sex-biased and unbiased genes on the X Chromosome and autosomes in *D. melanogaster*. Male-biased X Chromosome genes are flanked by regions with lower minimal HKAl, indicating that they are more likely to be evolving adaptively. B) Diverged genes located on the autosomes have significantly lower minimal HKAl within flanking regions than conserved genes. However, there is no clear pattern on the X chromosome. C) HKAl in flanking regions of diverged sex-biased and unbiased genes. Two-sided Mann-Whitney *U* test: *****p-value < 1e-5, ****p-value < 1e-4, ***p-value <

**Supplementary Figure S9.**
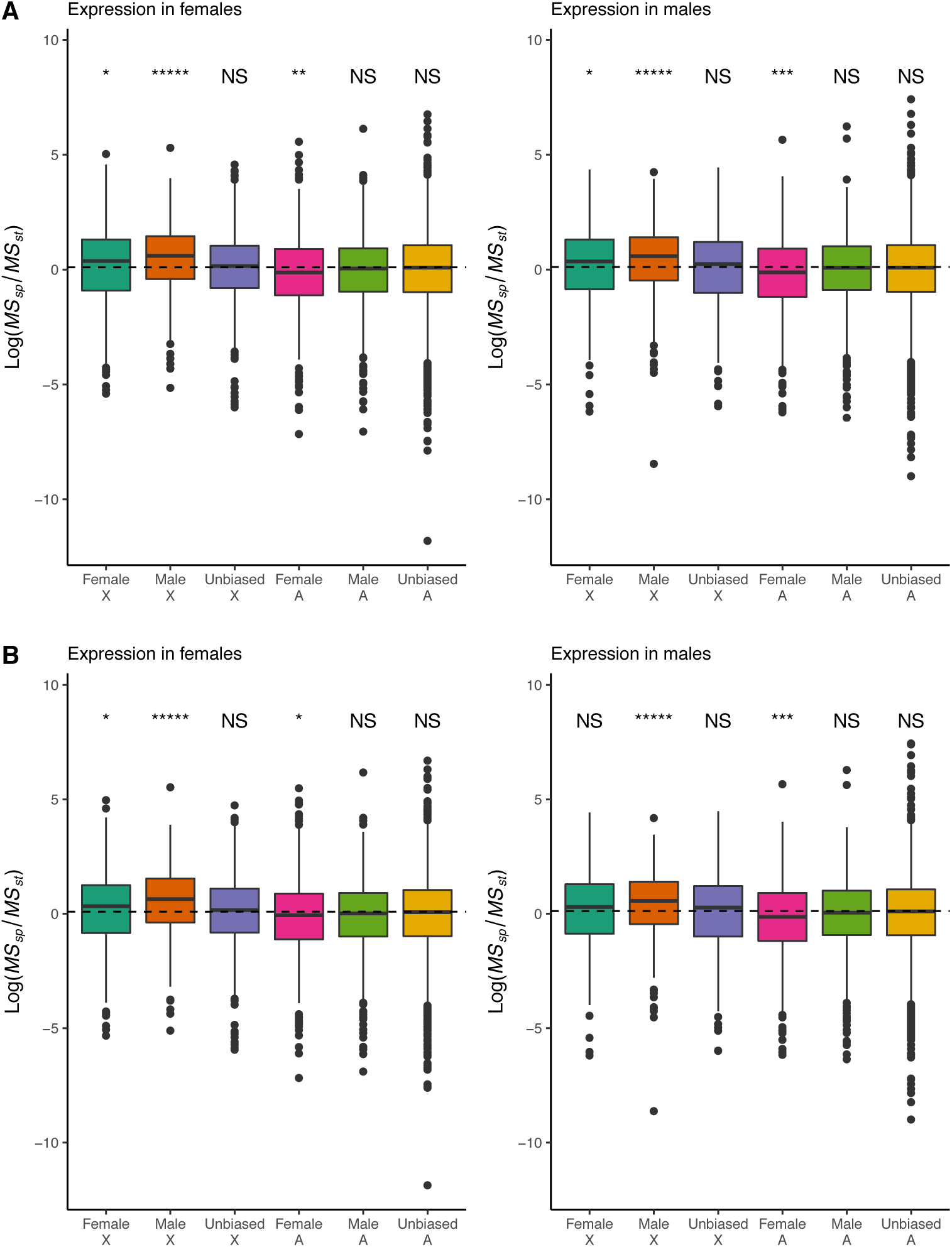
Ratio of interspecific to intraspecific variability in expression for genes in the *Drosophila* brain while controlling for tissue specificity. A) Ratio of interspecific to intraspecific variability while controlling for tissue specificity, with tissue specificity defined in *D. melanogaster*. B) Ratio of interspecific to intraspecific variability, with tissue specificity defined in *D. yakuba*. Sex-biased genes defined in the same way as in Figure 4.

**Supplementary Figure S10.**
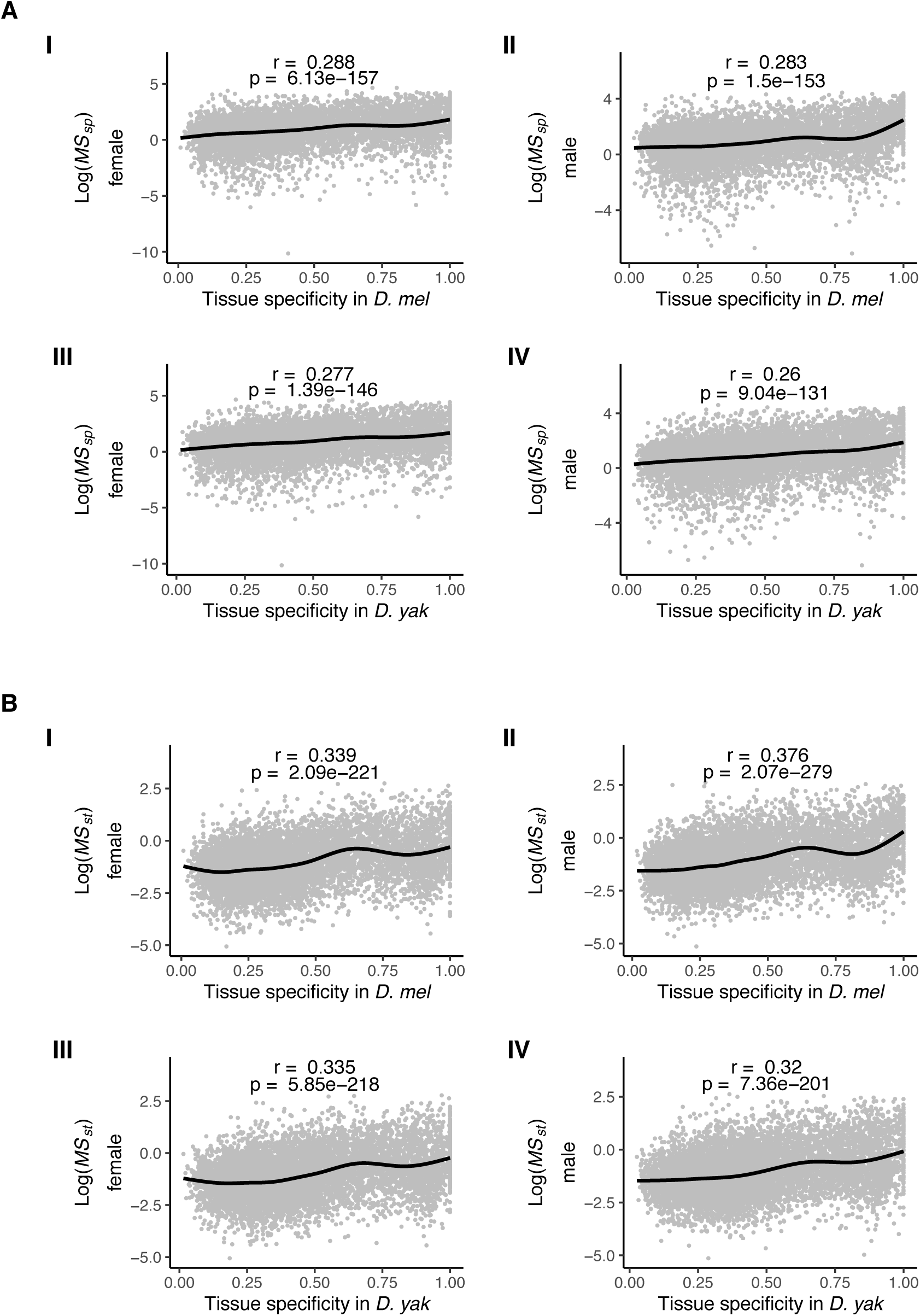
Positive relationship between gene expression variability and tissue specificity. A) Relationship between interspecific variability and tissue specificity in males and females for *D. melanogaster* (I and II) and *D. yakuba* (III and IV). B) Relationship between intraspecific variability and tissue specificity in males and females for *D. melanogaster* (I and II) and *D. yakuba* (III and IV). Correlations shown are Spearman correlations. Lines are generalized additive model smoothing curves. Sex-biased genes defined in the same way as in Figure 4.

**Supplementary Figure S11.**
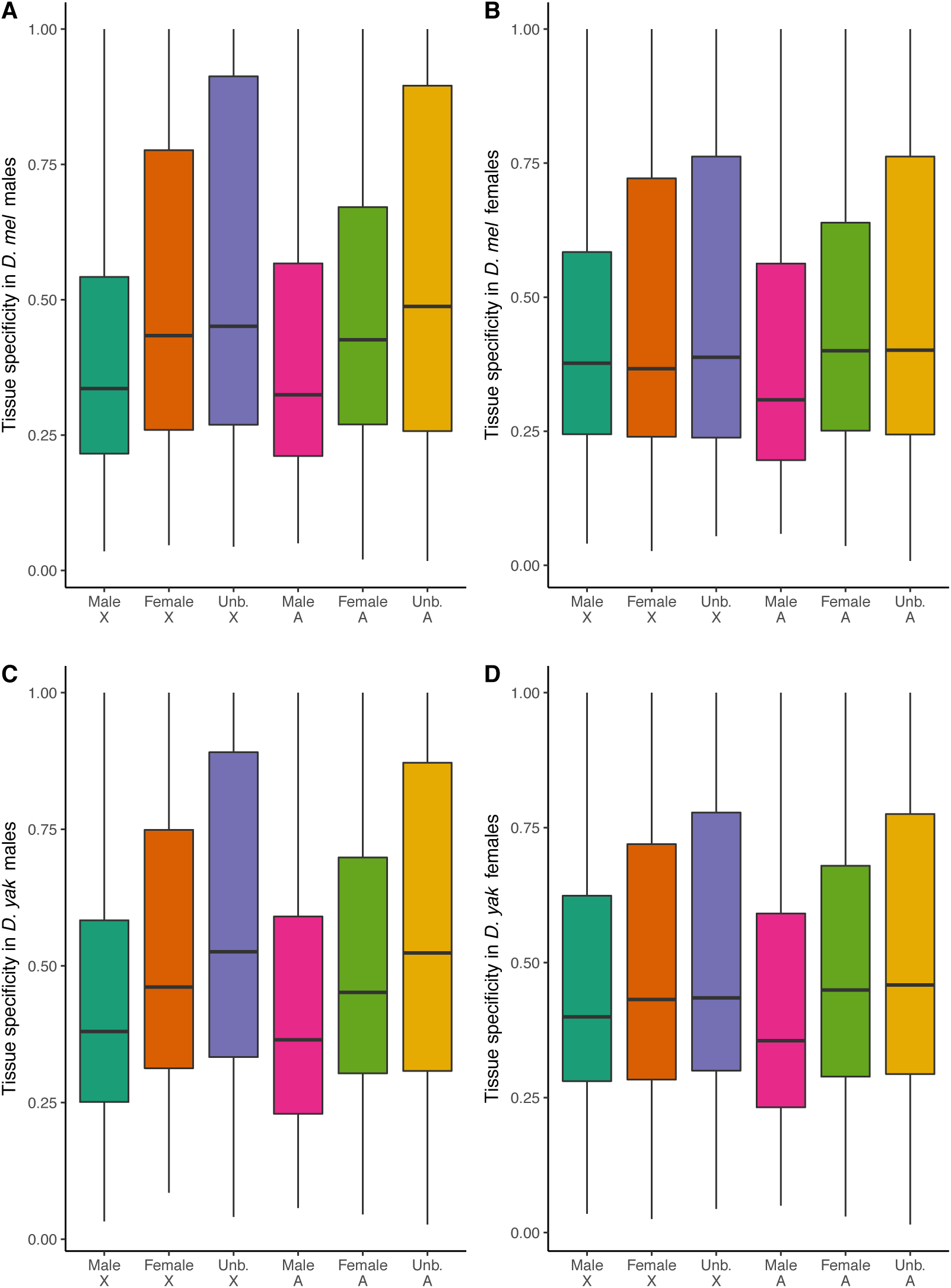
Tissue specificity of sex-biased genes. A) Tissue specificity in *D. melanogaster* males. B) Tissue specificity in *D. melanogaster* females. C) Tissue specificity in *D. yakuba* males. D) Tissue specificity in *D. yakuba* females. Sex-biased genes defined in the same way as in Figure 4.

**Supplementary Figure S12.**
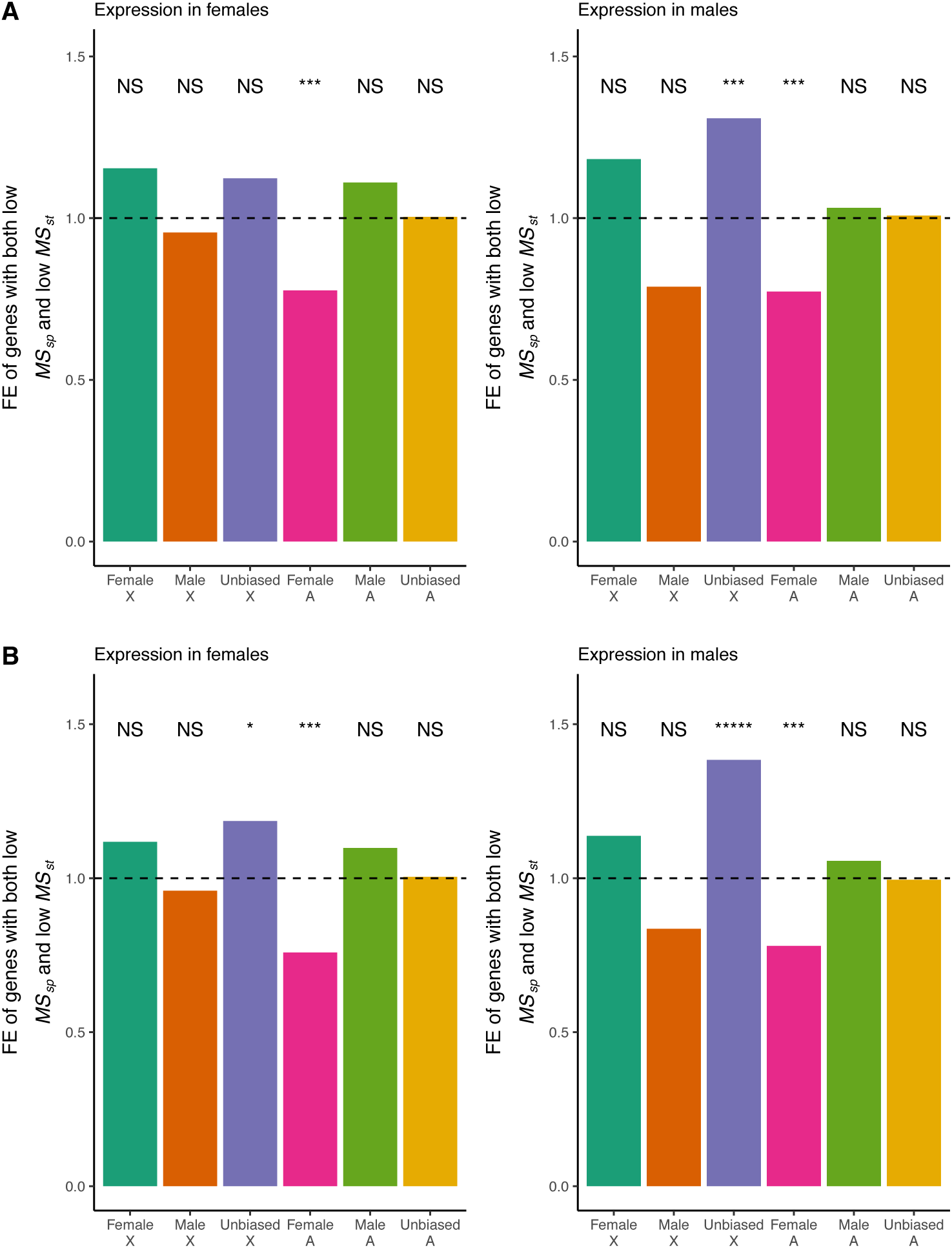
Fold enrichment of genes with both low intraspecific and low interspecific variabilities while controlling for tissue specificity. A) Controlling for tissue specificity in *D. melanogaster*. B) Controlling for tissue specificity in D. *yakuba*. Genes with low interspecific and intraspecific variabilities were defined as genes with *MS*_*species*_ and *MS*_*strain*_ both below the genome-wide medians. P-values were determined using a hypergeometric test followed by Benjamini-Hochberg correction. *****p-value < 1e-5, ****p-value < 1e-4, ***p-value < 1e-3, **p-value < 1e-2, *p-value < 5e-2. Sex-biased genes defined in the same way as in Figure 4.

**Supplementary Figure S13.**
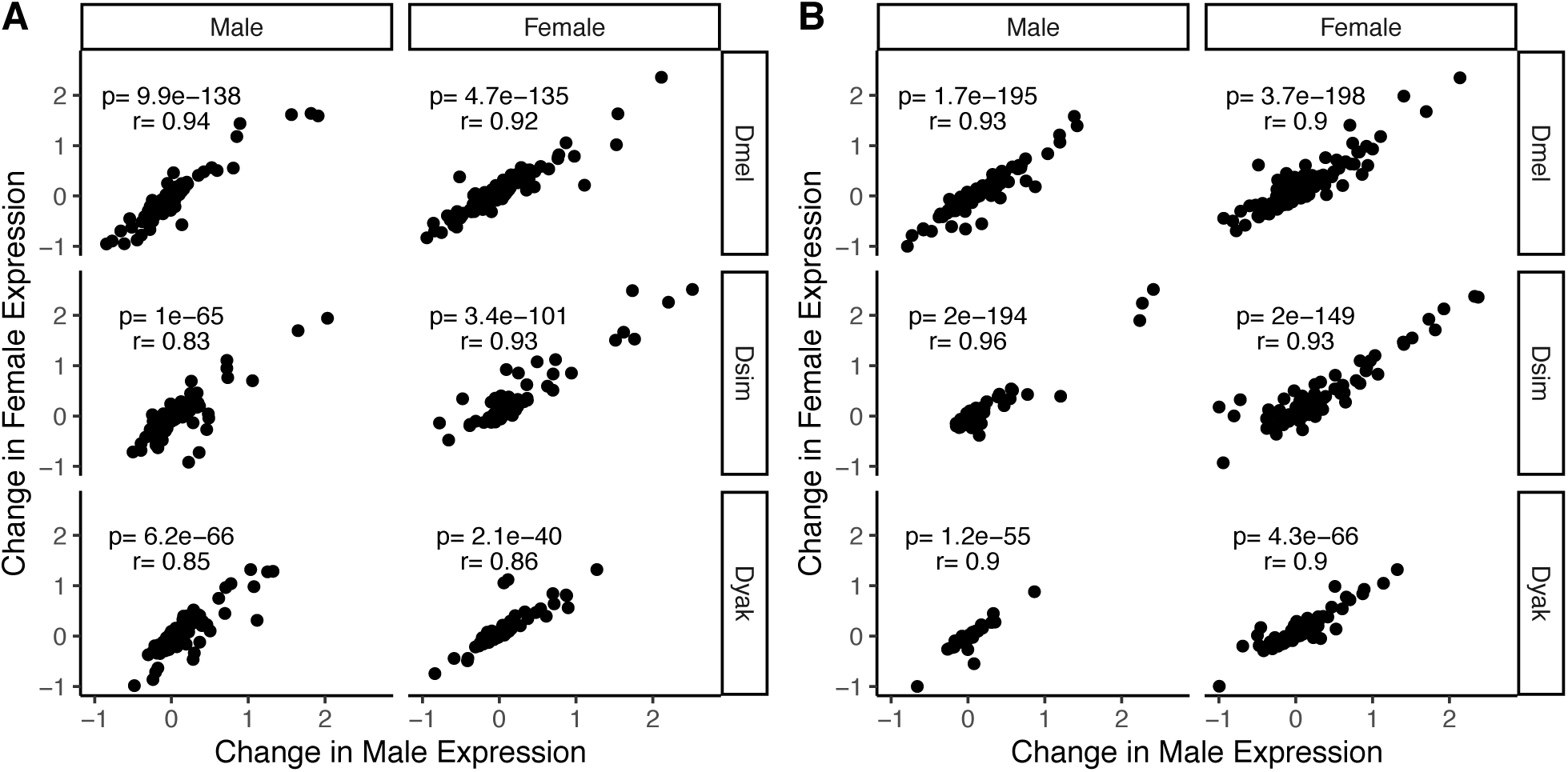
Change in expression relative to ancestral for state for sex-biased genes. A) The change in expression relative to ancestral state for sex-biased genes on the X Chromosome in the three *Drosophila* species. The ancestral expression levels were determined by maximizing the likelihood of a Brownian motion model on the phylogenetic tree. Ancestral states were calculated independently for males and females. The change in expression was defined as (current – ancestral)/abs(ancestral), where ‘current’ is the log_2_(TPM + 1) of the extant expression and ‘ancestral’ is the log_2_(TPM + 1) of the inferred ancestral expression. Then the Pearson correlation between changes in males and females were calculated. B) The change in expression relative to ancestral state for sex-biased genes located on the autosomes.

**Supplementary Figure S14.**
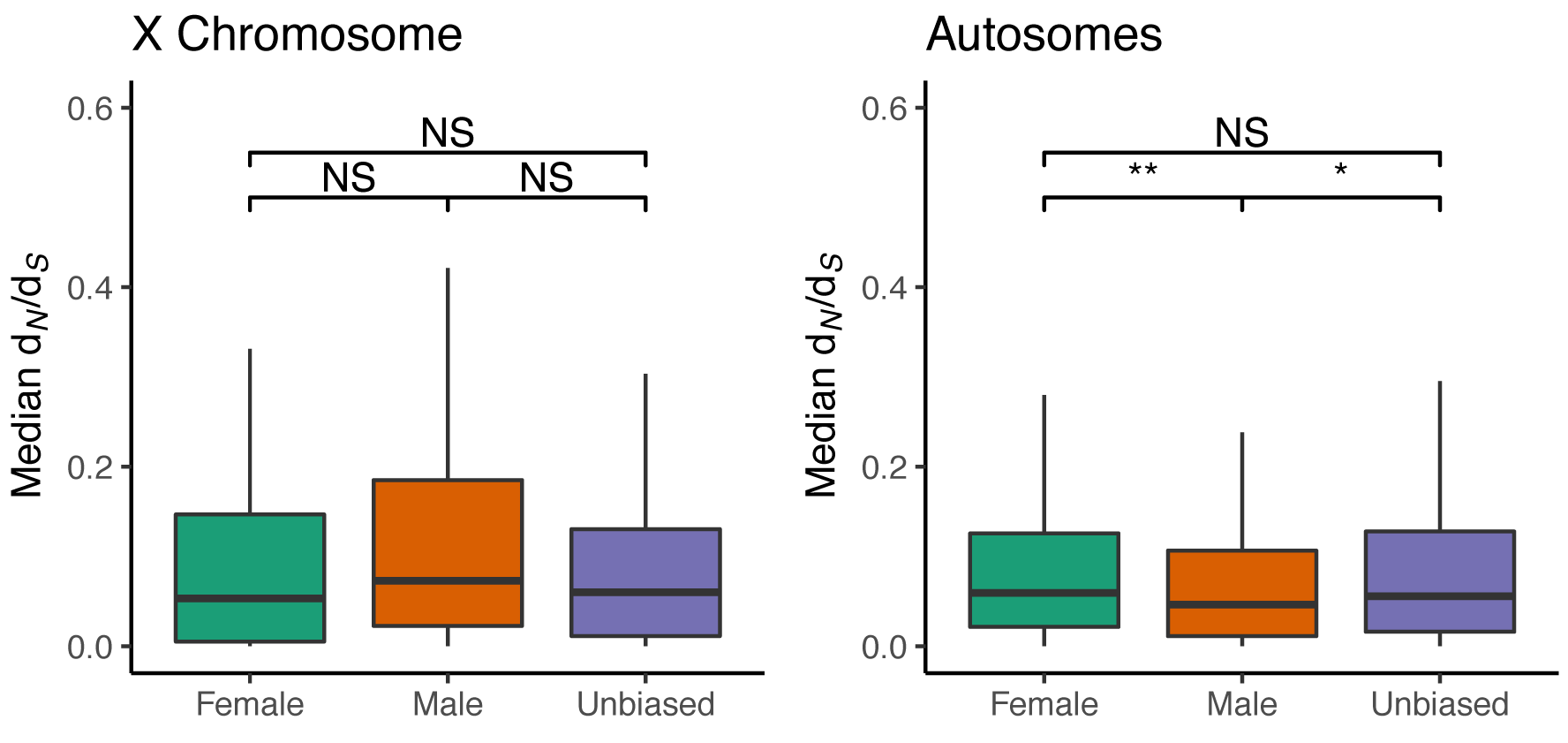
*d*_N_/*d*_S_ of sex-biased and unbiased genes. Male-biased autosomal genes have significantly lower *d*_N_/*d*_S_ than female-biased and unbiased autosomal genes. Two-sided Mann-Whitney *U* test with Benjamini-Hochberg correction: ***p-value < 1e-3, **p-value < 1e-2, *p-value < 5e-2. Sex-biased genes defined in *D. melanogaster* at a FDR of 0.05. Outliers were not included in the plots.

**Supplementary Table S5.** Test for faster-X expression evolution. Sex-biased genes located on the X Chromosome show a faster-X effect for gene expression evolution. *MS*_*species*_*/MS*_*strain*_ ratios were compared between genes located on the X Chromosome and all expressing genes located on the autosomes. Two-sided Mann-Whitney *U* test with Benjamini-Hochberg correction. Sex-biased genes defined in the same way as in Figure 4.

|  | p-Value<br>females | p-Value<br>males |
| --- | --- | --- |
| **Female-biased** | **6.16x10<sup>-3</sup>** | **6.69x10<sup>-3</sup>** |
| **Male-biased** | **1.07x10<sup>-11</sup>** | **2.00x10<sup>-9</sup>** |
| **Unbiased** | 7.90x10<sup>-2</sup> | 1.11x10<sup>-1</sup> |

**Supplementary Table S6.** McDonald-Kreitman test on sex-biased genes. A) X Chromosome sex-biased genes show an enrichment of genes (relative to all genes) with evidence of positive selection. B) When compared to genes on their native chromosomes there is no longer an enrichment. Sex-biased genes defined in *D. melanogaster* at a FDR of 0.05.

**A**
|  | Number of Genes<br>Tested | Number of<br>Significant Genes | Over/Under-Represented | p-Value<br>(Hypergeometric<br>Test.) |
| --- | --- | --- | --- | --- |
| X Chr. Male | 197 | 32 | Over | **2.94x10<sup>-5</sup>** |
| X Chr. Female | 200 | 32 | Over | **4.06x10<sup>-5</sup>** |
| Autosomal Male | 340 | 19 | Under | 1.54x10<sup>-1</sup> |
| Autosomal Female | 478 | 28 | Under | 1.31x10<sup>-1</sup> |

|  | p-Value (Hypergeometric<br>Test.) |
| --- | --- |
| X Chr. Male | 7.18x10<sup>-1</sup> |
| X Chr. Female | 7.97x10<sup>-1</sup> |
| Autosomal Male | 6.13x10<sup>-1</sup> |
| Autosomal Female | 6.87x10<sup>-1</sup> |

